# Developmental Cerebellar Pathology in Mouse Models of *SCN2A* Premature Termination Codon Variants

**DOI:** 10.64898/2026.09.13.751275

**Authors:** Katelin E.J Scott, Roy Gorter, Alayna Mull, Patrick Breheny, Chris A. Ahern, Aislinn J. Williams

## Abstract

Autism spectrum disorder is a neurodevelopmental disorder with both genetic and environmental contributors. *SCN2A*, the gene which encodes the alpha subunit of the voltage-gated sodium channel Na_v_1.2, is a known monogenetic risk factor for autism spectrum disorder. The cerebellum is frequently implicated in autism spectrum disorder and other neurodevelopmental disorders but has largely been unexplored in relation to *SCN2A* variants, especially from a developmental perspective. Na_v_1.2 is highly expressed within the cerebellum, specifically within cerebellar granule neurons, where it helps to drive action potential generation and propagation. Cerebellar granule neuron activity and maturation is crucial for shaping the development and morphology of the rest of the cerebellar cortex. Here we investigated early-postnatal cerebellar development in two mouse models carrying patient-derived *SCN2A* premature termination codon variants, *Scn2a-*p.Y84X and -p.R1627X. Overall, both *Scn2a* premature termination codon variant mouse lines displayed largely normal physical development and unaffected non-cerebellar developmental milestones. *Scn2a^Y84X/+^* mice, but not *Scn2a^R1627X/+^* mice, demonstrated alterations in cerebellar-driven motor behaviors, specifically faster performance in the surface righting reflex and cliff avoidance compared to wildtype littermates. Coinciding with the behavioral findings, *Scn2a* variants had divergent and age-dependent effects on cerebellar glutamatergic presynaptic marker expression, indicating that some, but not all, *Scn2a* premature termination codons impair glutamatergic synapse development. Both *Scn2a* variants exhibited changes to cerebellar cytoarchitecture, such as reductions in Purkinje cell density and soma size, which was more prominent in *Scn2a^Y84X/+^* mice, and faster migration of cerebellar granule neurons from the external granule layer to the internal, indicating a possible shift in the timing of cerebellar maturation. Our results situate the cerebellum as an early site for *SCN2A* pathophysiology and establish that cerebellar consequences of *SCN2A* premature termination codon variants may be position- and age-dependent, suggesting that influences beyond simple heterozygous loss of Nav1.2 drive phenotypes. Our characterization of how *Scn2a* premature termination codon variants differentially impair cerebellar development may help explain the heterogeneity of clinical presentations of *SCN2A* loss of function variants and potentially inform upon the timing of therapeutic intervention.

## Introduction

In the United States 1 in 6 children will be diagnosed with a developmental disability.^1^ One out of 31 of these patients will be diagnosed with autism spectrum disorder (ASD)^1^ by 8 years old. The etiology of ASD is complex, including genetics, early life experience, and the environment.^2^ However, some cases of ASD can be traced to a monogenic cause.^3^ One prominent monogenetic risk gene for ASD is *SCN2A,* which encodes the alpha subunit of Na_v_1.2, a voltage-gated sodium channel.^3^ Na_v_1.2 is involved in action potential generation and propagation for most glutamatergic neurons including cerebellar granule neurons (CGNs).^4–8^

Although ASD is heterogenous, it is frequently associated with molecular and functional changes in the cerebellum.^9–11^ For instance, reduced Purkinje cells (PC) size and density are common pathological findings in brains from ASD patients and rodent models.^9,10,12^ In rodent models of ASD, these changes in PCs can alter the ratio of inhibitory and excitatory cerebellar signals and change cerebellar output, leading to decreased adaptation and impaired accuracy in motor learning and movements.^9,13–15^ Furthermore, the cerebellum has notable roles in social and cognitive function, which are important in neurodevelopmental disorders like ASD.^9,16^

Na_V_1.2 is highly expressed in CGNs,^17–19^ where it is localized along axons called parallel fibers (PF) and axon terminals, which synapse onto PCs.^19^ CGNs provide direct excitatory input to PCs, the sole output of the cerebellar cortex.^20^ Through direct excitation of PC dendrites, CGNs play an integral role in regulating PC activity.^20–22^ These connections are particularly important during early development, when CGN input to PCs refines cerebellar cytoarchitecture and the interactions between both cell types reciprocally promotes maturation of each.^21–23^

Although Na_v_1.2 is highly expressed within the cerebellum, the vestibular ocular reflex (VOR) remains the only rigorously evaluated cerebellum-dependent behavior in any *Scn2a* model.^3,24^ Adult *Scn2a^+/-^* mice exhibit increased VOR (the ratio of the eye velocity to head velocity) in a gain-down protocol compared to wildtype controls.^24^ This was also observed in human patients carrying *SCN2A* loss of function (LOF) variants, which reduce Na_v_1.2 sodium current through impaired protein trafficking and/or protein truncation.^24^ While altered VOR indicates cerebellar involvement in *SCN2A* LOF, these findings do not establish when cerebellar dysfunction emerges or its underlying developmental mechanisms.

While previous studies indicate that Na_v_1.2 may be important for cerebellar-driven behavior, this has not been explored during early-postnatal development, nor in models containing human patient mutations. To address this gap, we examined early-postnatal cerebellar structural development and cerebellar-related behaviors in two mouse models carrying patient-derived premature termination codon variants (PTCs), *Scn2a-*p.Tyr84X (Y84X) and - p.Arg1627X (R1627X).^25^ These PTC variants reduce Na_v_1.2 protein expression to approximately 50% of wildtype.^25^ We assessed the effects of each variant on early cerebellar development through a battery of behavioral assays and histology techniques. Our results reveal that the cerebellum is a region of early pathophysiology for *SCN2A* PTC mutations, but cerebellar phenotypes are divergent across PTCs.

## Materials and Methods

### Animals

The generation of *Scn2a^Y84X/+^*and *Scn2a^R1627X/+^* mice was previously described.^25^ All experiments were performed in accordance with the National Institutes of Health guidelines for animal care and were approved by the University of Iowa’s Institutional Animal Care and Use Committee.

### Developmental Behavioral and Health Assessments

Assay methods were adapted from the literature.^26–28^ Mice were housed under a regular light cycle with lights on/off at 0900/2100 DST (0800/2000 non-DST). For behavioral studies, breeding cages used sires carrying the mutation and wildtype dams to minimize potential rearing differences between wildtype and heterozygous dams. *Scn2a^Y84X/+^* and *Scn2a^R1627X/+^*mice and their respective wildtype littermates were tested daily during the light cycle from postnatal day (PND) 1 to PND12. Pups were identified on PND1 by hind toe clip. Pups were evaluated in the order they were selected from the home cage with ≥5 minutes between assays. PND1 pups were only evaluated for weight. PND2 pups were evaluated for weight, eye opening, pinna detachment, forelimb grasp, surface righting, and cliff avoidance. From PND3 onward, negative geotaxis was added after cliff avoidance. Researchers remained blinded to genotype until statistical analysis. Six to eight animals were used per sex and genotype group.

#### Health Measurements

Pups were weighed daily from PND1 to PND12. Eye opening and pinna detachment were monitored from PND1 to until occurrence and were scored as follows: 0 = no or 1 = yes.

#### Forelimb Grasping Reflex

Forelimb grasping reflex was tested daily from PND2 to PND12 using a straightened paperclip held 1in from a padded surface. Forelimb grasping reflex was scored as follows: 0 = no grasp, 1 = paws folded over paperclip, 2 = able to hang from paperclip for ≥ 1 second.□

#### Surface Righting Reflex (SRR)

SRR was tested daily from PND2 to PND12. Pups were placed in a supine position and recorded for 30 seconds or until the pup righted itself with all four paws on the ground, whichever occurred first. SRR was scored as follows: 0 = unsuccessful, 1 = successful. The latency for each pup to right their body from a supine position to a prone position with all four paws on the ground was recorded.

#### Cliff Avoidance (CA)

CA was tested daily from PND2 to PND12. A solid flat platform made from a clipboard wrapped in a bench pad was placed perpendicularly over an open standard housing cage to create a “cliff”. Bedding was added to the cage to reduce falling distance to 10cm or less. Pups were placed with their nose and forepaws wrapped just over the edge of the platform facing towards the bedding. Pups were observed for 30 seconds or until they turned 180° away from the “cliff”. CA was scored as follows: 0 = fall, 1 = no movement, 2 = partial turn <180° from the edge, 3 = full 180° turn away from edge. If the mouse scored a 3, then latency to turn was also recorded.

#### Negative Geotaxis (NG)

Negative geotaxis was tested daily from PND3 to PND12. Pups were placed on a negative slope of 45° created from a clipboard wrapped in a bench pad with their nose pointed downward. The pups were evaluated for 60 seconds or until they turned 180° orienting their body up the slope. The assay was scored as follows: 0 = fall off/stay/move downwards/turn less than 180°; 1 = turn their body 180° so their nose pointed upward. If the mouse scored a 1, the latency was also recorded.

#### Ultrasonic Vocalizations (USV)

Litters used for USV recordings were separate from those used for other behavioral measurements. For USV recordings, individual PND6 pups were isolated from their dam and placed into a glass slide staining dish (interior dimensions: 95 mm × 76 mm × 64 mm) with fresh bedding inside a larger styrofoam box within a sound attenuating chamber (MedAssociates, Cat: ENV-018). A microphone (Avisoft Bioacoustics, Cat: CM16/CMPA) was embedded into the box lid 5cm above the pup. Pups were recorded for 5 minutes using the Avisoft Bioacoustics UltraSoundGate 116H ultrasound recording interface and Avisoft-SASLab Pro recording software (Avisoft Bioacoustics). Data were analyzed using the LMT USV toolbox.^29^

### Histology and Microscopy

#### Tissue Processing

Methods were adapted from Hayes et al. and Klomp et al.^30,31^ Brain tissue was examined histologically at PND6, PND12, and PND21. Mice > PND6 were anesthetized with 17.5mg/ml Ketamine/2.5mg/ml Xylazine (0.1ml/20g) and transcardially perfused with 0.1M phosphate buffer (PB), followed by 4% paraformaldehyde in 0.1M PB. Whole brains were then dissected, immersed in 30% sucrose for ≥ 24 hours, rinsed in 0.1M PB and frozen in Optimal Cutting Temperature compound (Fisher Scientific Cat: 23-730-571) using a dry ice/2-methylbutane slurry. PND6 mice were euthanized by decapitation. Brains were dissected, rinsed in 0.1M PB, immersion-fixed in 4% paraformaldehyde in 0.1M PB for ≥ 24 hours, and processed as above. Tissues were stored at −80°C until sectioning. Brain tissue was sectioned sagittally using a cryostat at 20μm. Serial cerebellar vermis sections were used for immunofluorescence studies with every 5th section retained for Nissl-based experiments. Sections were air-dried and stored at −20°C until use.

Immunostaining:

Methods were adapted from Klomp et al.^30^ Nonspecific labeling was blocked for ≥ 4hr with blocking buffer (5% normal donkey serum in 0.01% Triton-X in phosphate buffered saline). All antibodies were diluted in blocking buffer. Primary antibodies included Vglut1 (1:200; Synaptic Systems, Cat:135-302)^21^, Vglut2 (1:200; Synaptic Systems, Cat: 135-403)^21^ and Calbindin D28K (1:200; Sigma Cat: C9848).^32^ Sections were incubated in primary antibodies overnight at 4°C. Secondary antibodies included Donkey α Rabbit 594 (1:500; Jackson, Cat:711-585-152)^31^ and Donkey α Mouse 488 (1:500; Jackson, Cat: 715-545-151).^31^ Nuclei were labeled with DAPI (1:1000; Thermo Scientific, Cat: D1306)^31^. Sections were incubated in secondary antibodies at room temperature for 4 hours. Coverslips were mounted in Mowiol (Sigma Aldrich, Cat:81381) containing 2.5% 1,4-diazobicyclo-[2.2.2]-octane (DABCO, Sigma, Cat: D2522).^33^

#### Nissl Stain

Methods were described in Hayes et al.^31^ Briefly, 20μm sagittal cerebellar vermis sections approximately 100μm apart were stained with thionin. Sections were rehydrated in deionized (DI) H_2_O for 5 seconds, followed by approximately 90 seconds in thionin stain with a quick rinse in DI H_2_O. Sections were submerged in 70% ethanol for 5 minutes followed by another 5 minutes in 95% ethanol, then submerged in a 95% alcohol/acetic acid solution until the desired intensity was achieved. Slides were then rinsed in 100% ethanol twice for 5 minutes each and cleared three times in Citrisolv (VWR International Incorporated, Cat: 5989-27-5) for 5 minutes each. Coverslips were mounted with Permount

#### Microscopy

Methods are as described in Klomp et al.^30^ Sections were imaged on an Olympus IX83 fluorescence microscope using Olympus CellSens Dimension 2.3 software. Analysis was done in ImageJ. For microscopy experiments, 6 to 8 animals per genotype/age group were used for analysis. Datasets were equally balanced between male and female animals with *n* = 3-4 per sex/genotype/age group.

### Image Analysis

#### Vglut Density

Methods were adapted from van der Heijden et al.^21^ Serial sagittal cerebellar vermis sections stained with anti-Vglut1 or anti-Vglut2 and anti-Calbindin were used for quantification. Images were taken at 40X magnification. The polygon tool in ImageJ was used to define the region of interest (ROI) around calbindin positive areas (PC somata and dendrites).

Background was removed with the subtract background tool (rolling ball radius: 3.6 pixels). Brightness and contrast were adjusted to visualize individual puncta. Images were thresholded using the threshold function. The analyze particles tool was used to quantify the percentage of each ROI that was positive for Vglut1 or Vglut2. Data from 2-3 images per lobule were averaged to generate one percentage value per ROI (*n* = 6–8 per genotype, *n* = 3–4 per sex/lobule).□

#### PC Density and Soma Size

Methods were adapted from Hayes et al. ^31^ Images were analyzed in ImageJ. Scale was calibrated by measuring the length of the scale bar (known distance) to convert distance in pixels to microns. For density, two thionin-stained sagittal cerebellar vermis sections 100µm apart (*n* = 6–8 per genotype, *n* = 3–4 per sex) were imaged at 20X. The straight-line tool was used to draw a line across the PC monolayer spanning 3–10 consecutive PCs (Supplemental Fig. 1). Full PC somas intersected by the line were counted and line length was recorded. Two to four measurements were taken from 4 consistent regions in larger lobules (anterior base, posterior base, anterior crown, posterior crown) and from 2 regions (anterior and posterior) for smaller lobules (VII and X; Supplemental Fig. 1). . Values were averaged to calculate the PC density for each lobule per one hundred micrometers (density = (number of cells / lengths of line) *100). For soma area, one thionin-stained sagittal cerebellar vermis section (*n* = 6-8 per genotype, *n* = 3-4 per sex) was imaged at 20X magnification. Using the polygon tool, PC somata with visible primary proximal dendrites (2-8 per section) were traced from lobules as described for PC density. These values were averaged together to generate a single data point per mouse/lobule.

#### Cerebellar Cortex Layer Thickness

Sections stained with DAPI (PND6) or thionin (PND12 and PND21) were used for this quantification. Images from sagittal cerebellar vermis sections of the external granule layer at PND6 and PND12 and the molecular layer at PND12 and PND21 were taken at 20X magnification and analyzed in ImageJ. The image scale was calibrated as described above. One measurement was taken from each of the regions described in the methods for PC soma area using the straight-line tool (Supplemental Fig. 1). Individual measurements were averaged to generate a single data point per mouse/lobule (*n* = 6-8 per genotype/age, *n* = 3-4 per sex/genotype/age).□

### Protein Quantification

#### Capillary immunoassay

Methods were modified from Al Saneh and Hermosillo Arrieta et al.^25^ Mice were euthanized at PND21 via isoflurane followed by decapitation after toe-pinch reflex was undetectable. Brains were rapidly extracted and cerebelli were isolated from the cortex and brainstem by cutting just below the cerebellar peduncles. Cerebelli were bisected along the midsagittal plane through the vermis. Half cerebellum was homogenized in RIPA lysis buffer (Thermo Fisher Scientific, Cat: PI89901) supplemented with cOmplete™ Mini EDTA-free protease inhibitor (Roche, Cat:11836170001). Protein concentration was determined using a Bradford assay (Bio-Rad, Cat: 5000002). The Jess™ Simple Western™ capillary immunoassay system was used to quantify Vglut1 and Vglut2 protein expression using the total protein assay for normalization (Bio-Techne, Cat: DM-TP01). Protein lysate was loaded at 0.026 mg/mL for Vglut1 and 0.04 mg/mL for Vglut2. The Vglut1 antibody (described above) was used at 1:1000 while the Vglut2 antibody (described above) was used at 1:500.

### Statistical Methods

All statistical analyses and graphical representations were conducted using R Studio (R 4.5.1). Early reflex behaviors were analyzed from the first day any animal fully performed the reflex to the day all animals completed the reflex at least once. No animals or trials were excluded from analysis. Data were analyzed using linear mixed effects modeling (LMEM) in R (R 4.1.1, emmeans 1.7.0, lme4 1.1.27.1, lmerTest 3.1-3, effectsize 0.5). Individual mouse identity nested within litter was included as a random effect.^34^ For USV and day of occurrence, analysis was completed with litter as a random effect without mouse identity nested. Estimated marginal means (EMM) were used to follow up significant interaction effects. Results were considered significant when p□≤□0.05 (denoted in all graphs as follows: *p□≤□0.05; **p□≤□0.01).

## Results

### General health measurements are not altered by *Scn2a* PTCs

We explored baseline health characteristics of heterozygous *Scn2a* PTC mice during early-postnatal development. From PND1 to PND12, neither *Scn2a^Y84X/+^* or *Scn2a^R1627X/+^* mice demonstrated differences in weight or the age at which they first opened their eyes compared to their respective wildtype littermates (Fig. 1A-D; Supplemental Table 1). *Scn2a^Y84X/+^* did not demonstrate a difference in pinna detachment, however, female *Scn2a^R1627X/+^* mice underwent complete pinna detachment slightly later than all other groups (LMEM: wildtype v. R1627X genotype*sex, F(D_1_,D_32.71_) = 5.19, *p* = 0.03; EMM: females, *t*(9.07) = −2.09, *p*□= 0.05; Fig. 1E,F; Supplemental Table 1). We do not expect that the slowed rate of pinna detachment in female *Scn2a^R1627X/+^* mice impeded the animals’ ability to perform behavioral assays.

**Figure 1:**
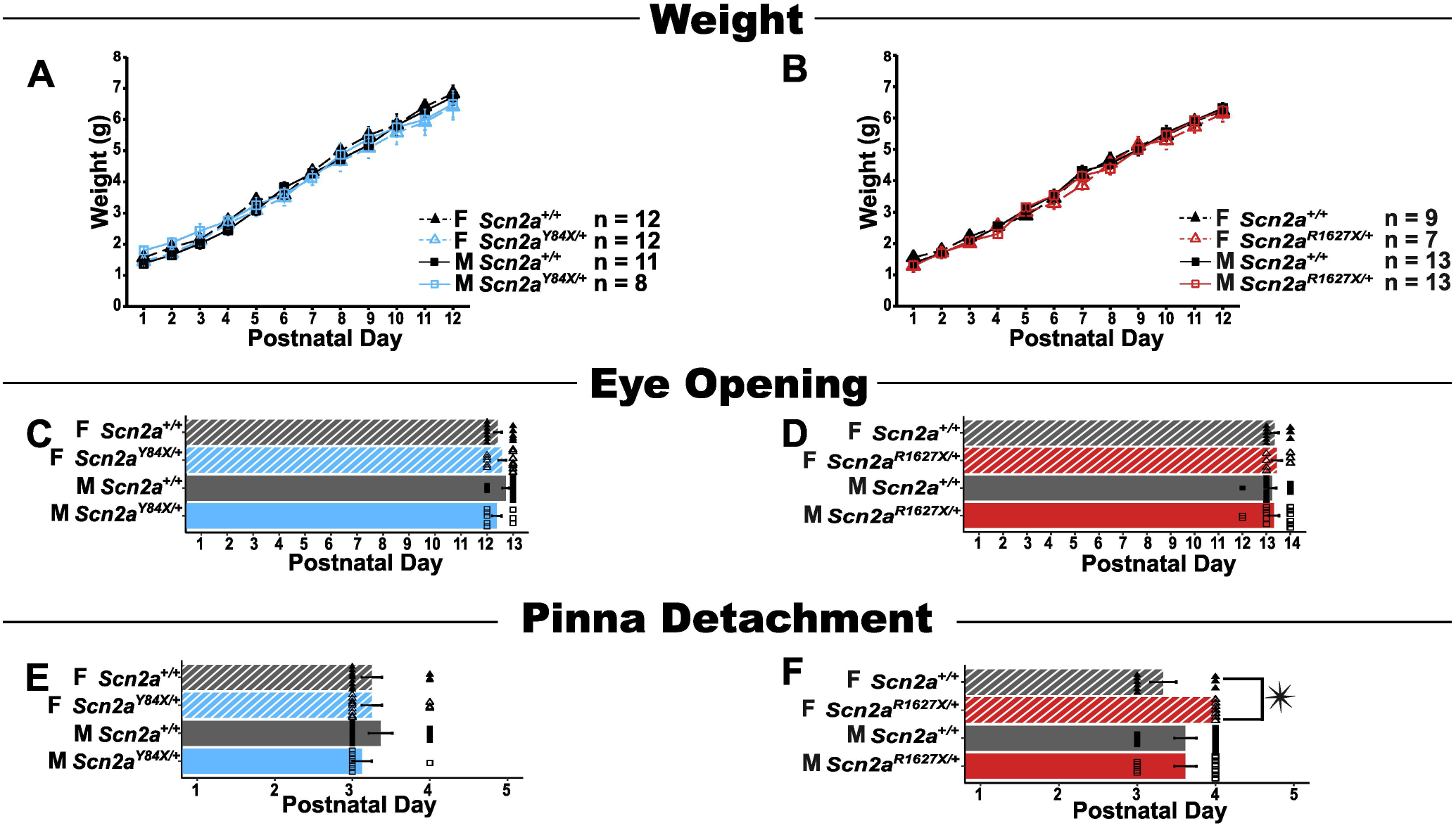
Mutant *Scn2a* PTC mice have largely normal physical development. **A)** Weight of *Scn2a^Y84X/+^* pups and **B)** *Scn2a^R1627X/+^* pups were similar to wildtype littermates (LMEM: wildtype v. Y84X, genotype, F(D_1_,D_38.51_) = 0.19, *p* = 0.66; LMEM: wildtype v. R1627X, genotype, F(D_1_,D_33.69_) = 1.94, *p* = 0.17). **C)** The day when *Scn2a^Y84X/+^* and **D)** *Scn2a^R1627X/+^* mouse pups fully opened eyes was similar to respective wildtype littermates (LMEM: wildtype v. Y84X, genotype, F(D_1_,D_35.74_) = 0.51, p = 0.48; LMEM: wildtype v. R1627X, genotype, F(D_1_,D_34.83_) = 0.05, *p* = 0.83). **E)** The day when *Scn2a^Y84X/+^* mouse pup pinnae fully detached from skull was similar to wildtype littermates (LMEM: wildtype v. Y84X, genotype, F(D_1_,D_33.97_) = 0.01, *p* = 0.95). **F)** The day when *Scn2a^R1627X/+^* female mouse pup pinnae fully detached from skull was slightly delayed compared to female wildtype littermates (LMEM: wildtype v. R1627X, genotype*gex, F(D_1_,D_32.71_) = 5.19, p = 0.03; EMM: females, *t*(9.07) = −2.09, *p*□= 0.05). Data are represented as mean +/- SEM. (PTC = premature termination codon, LMEM = linear mixed effects model, EMM = estimated marginal means, PND = postnatal day)

### Cerebellar-driven early reflexes are altered in *Scn2a^Y84X/+^* mice

Since there is evidence that Na_V_1.2 deficiency in adulthood causes cerebellar-related phenotypes,^24^ we evaluated early cerebellar-related reflexes in *Scn2a^Y84X/+^* and *Scn2a^R1627X/+^* mice. We hypothesized that Na_V_1.2 deficiency would impair development of early reflex behaviors due to Na_V_1.2’s role in the generation and propagation of action potentials, its importance in CGN activation of PCs, and the subsequent long-term potentiation between the two cell types.^7,19,24^ *Scn2a^Y84X/+^*mice were quicker to perform the cliff avoidance (CA) reflex than wildtype (LMEM: wildtype v. Y84X, genotype, F(D_1_,D_37.24_) = 6.84, *p* = 0.01). A genotype by day interaction revealed their faster performance on PND8 and PND9 (LMEM: wildtype v. Y84X, day*genotype, F(D_8_,D_311.22_) = 2.93, *p* = 0.004; EMM: PND8, *t*(340) = 2.62, *p*□= 0.009; EMM:

PND9, *t*(340) = 4.62, *p*□= <.0001). *Scn2a^R1627X/+^*mice, however, performed similarly to wildtype on CA (Fig. 2A,B; Supplemental Table 2). Both *Scn2a^Y84X/+^* and *Scn2a^R1627X/+^* mice demonstrated the same skill level and first performed CA at a similar age to their respective wildtype littermates (Fig. 2C-F; Supplemental Table 2). Likewise, *Scn2a^Y84X/+^* mice and male mice were quicker to perform the surface righting reflex (SRR) than wildtype and female littermates (LMEM: wildtype v. Y84X, genotype, F(D_1_,D_39_) = 4.79, *p* = 0.04; sex, F(D_1_,D_39_) = 4.58, *p* = 0.04). Male *Scn2a^Y84X/+^* mice, when compared to male wildtype, were faster on the SRR (LMEM: wildtype v. Y84X, genotype*sex, F(D_1_,D_39_) = 6.47, *p* = 0.02; EMM: males, *t*(37.2) = 2.93, *p*□= 0.006), but this effect was not seen for female *Scn2a^Y84X/+^* mice nor *Scn2a^R1627X/+^* mice (Fig. 2G, H; Supplemental Table 3). Male *Scn2a^Y84X/+^* mice also had higher skill performance scores than male wildtype mice on the SRR (LMEM: wildtype v. Y84X, genotype*sex, F(D_1_,D_39_) = 4.33, *p* = 0.04; EMM: males, *t*(37.2) = −2.13, *p*□= 0.04), but neither female *Scn2a^Y84X/+^* nor *Scn2a^R1627X/+^* mice outscored wildtype (Fig. 2I, J; Supplemental Table 3). Both *Scn2a^Y84X/+^* and *Scn2a^R1627X/+^* mice first performed the SRR at the same age as their respective wildtype littermates (Fig. 2K,L; Supplemental Table 3). No differences in performance or first day of achievement on negative geotaxis were identified for *Scn2a^Y84X/+^* or *Scn2a^R1627X/+^* mice (Supplemental Fig. 2A-F; Supplemental Table 4).

**Figure 2:**
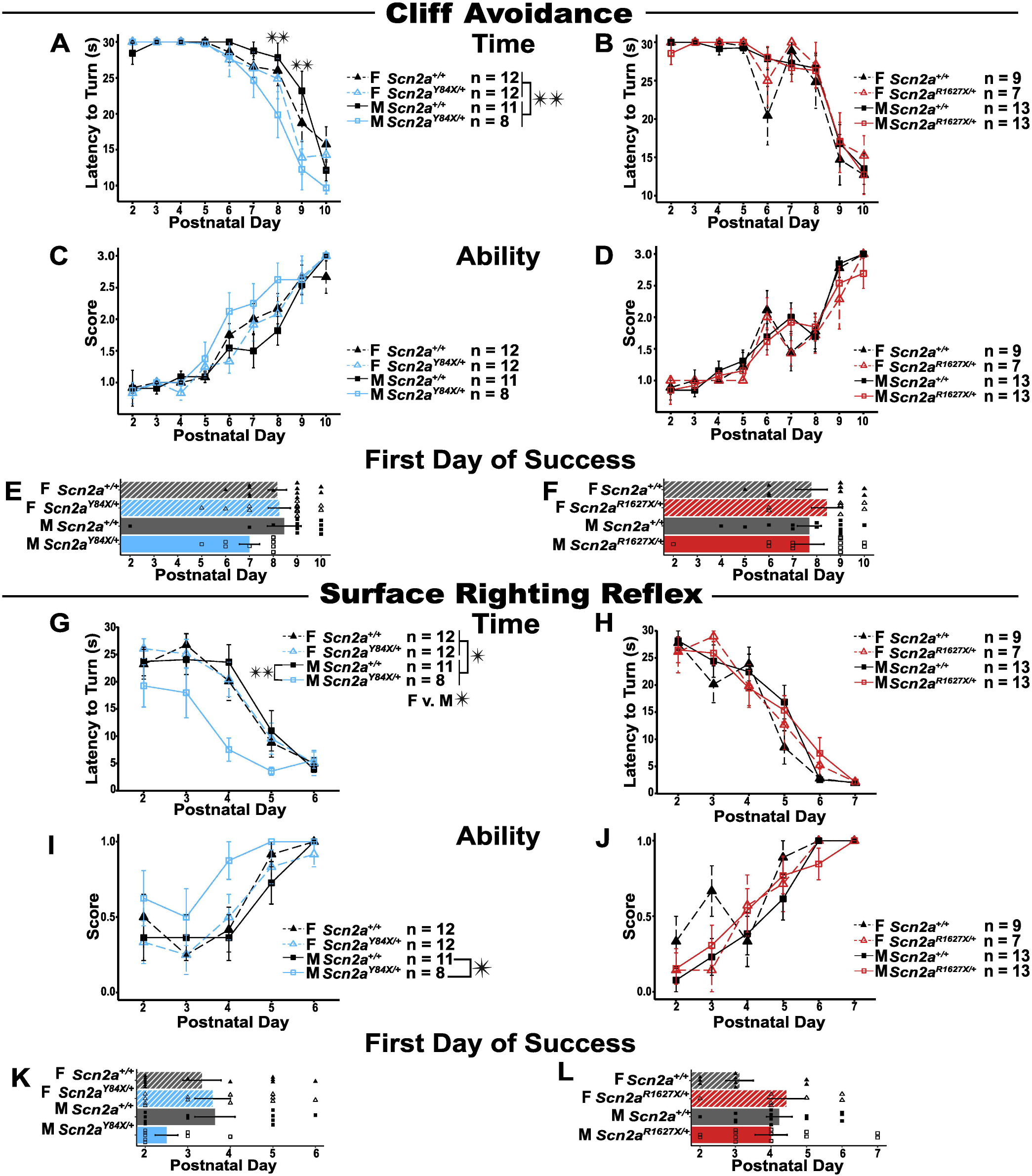
*Scn2a^Y84X/+^* mice display some cerebellar related behaviors earlier than wildtype littermates. **A)** *Scn2a^Y84X/+^* mice are faster than wildtype littermates in cliff avoidance (LMEM: wildtype v. Y84X, genotype, F(D_1_,D_37.24_) = 6.84, *p* = 0.01). This difference was most apparent on PND8 and PND9 (LMEM: wildtype v. Y84X, genotype*day, F(D_8_,D_311.22_) = 2.93, p = 0.004; EMM: PND 8, t(340) = 2.62, *p* = 0.01; EMM: PND9, t(340) = 4.62, *p* =<.0001). **B)** *Scn2a^R1627X/+^* mice are similar to wildtype littermates in cliff avoidance (LMEM: Wildtype v. R1627X, genotype, F(D_1_,D_37.42_) = 0.19, *p* = 0.67) **C)** *Scn2a^Y84X/+^* mice and **D)** *Scn2a^R1627X/+^* mice score similarly to their respective wildtype littermates on cliff avoidance (LMEM: wildtype v. Y84X, genotype, F(D_1_,D_38.52_) = 1.49, *p* = 0.23; LMEM: Wildtype v. R1627X, genotype, F(D_1_,D_310.13_) = 0.23, *p* = 0.63). **E)** *Scn2a^Y84X/+^* mice and **F)** *Scn2a^R1627X/+^* mice first achieve cliff avoidance on a similar day to their respective wildtype littermates (LMEM: wildtype v. Y84X, genotype, F(D_1_,D_38.99_) = 1.64, *p* = 0.21; LMEM: wildtype v. R1627X, genotype, F(D_1_,D_37.58_) = 0.05, *p* = 0.82). **G)** *Scn2a^Y84X/+^*mice and male mice are faster than wildtype and female littermate littermates respectively on surface righting reflex (LMEM: wildtype v. Y84X, genotype, F(D_1_,D_39_) = 4.79; LMEM: sex, F(D_1_,D_39_) = 4.58, *p* = 0.04). Specifically, *Scn2a^Y84X/+^* male mice are much faster than male wildtype littermates (LMEM: wildtype v. Y84X, genotype*sex, F(D_1_,D_39_) = 6.47, p = 0.02; EMM: males, t(37.20) = 2.93, *p* = 0.006). **H)** *Scn2a^R1627X/+^* mice speed on surface righting reflex is similar to wildtype littermates (LMEM: wildtype v. R1627X, genotype, F(D_1_,D_227.32_) = 0.28, *p* = 0.60). **I)** *Scn2a^Y84X/^*^+^ male mice score better than wildtype littermates on surface righting reflex (LMEM: wildtype v. Y84X, genotype*sex, F(D_1_,D_39_) = 4.33, *p* = 0.04; EMM: males, t(37.20) = −2.13, *p*□= 0.04). **J)** *Scn2a^R1627X/+^* mice surface righting reflex score is similar to wildtype littermates (LMEM: wildtype v. R1627X, genotype, F(D_1_,D_35.05_) = 0.12, *p* = 0.73). **K)** *Scn2a^Y84X/+^* mice and **L)** *Scn2a^R1627X/+^* mice first achieve the surface righting reflex on a similar day to their respective wildtype littermates (LMEM: wildtype v. Y84X, genotype, F(D_1_,D_38.98_) = 1.02, *p* = 0.32; LMEM: wildtype v. R1627X, genotype, F(D_1_,D_37.83_) = 1.22, *p* = 0.28). Data are represented as mean +/- SEM. (LMEM = linear mixed effects model, EMM = estimated marginal means)

### Non-cerebellar driven early reflex behaviors are unchanged in *Scn2a* PTCs

We also examined non-cerebellar driven behaviors in *Scn2a^Y84X/+^* and *Scn2a^R1627X/+^* mice during early-postnatal development by evaluating the forelimb grasping reflex and observing USVs. Neither *Scn2a^Y84X/+^* or *Scn2a^R1627X/+^* mice exhibit differences in skill level nor in the age at which they first achieved the forelimb grasping reflex (Supplemental Fig. 3A-D; Supplemental Table 5). We also observed no differences in the count, duration, frequency, or number of modulations in early-postnatal USVs in *Scn2a^Y84X/+^* or *Scn2a^R1627X/+^* mice compared to their respective wildtype littermates (Supplemental Fig. 4A-H; Supplemental Table 6).

### Differential impacts of *Scn2a* PTCs on cerebellar excitatory presynaptic markers

CGNs are the predominant source of cerebellar Vglut1 expression, which is enriched at presynaptic sites of excitatory synapses.^19^ We hypothesized that reduced Na_V_1.2 expression in *Scn2a* PTC mice would impair PF excitatory synapse formation, resulting in decreased Vglut1 expression during early-postnatal development. We therefore examined Vglut1 puncta distribution in the PC layer and molecular layer of the cerebellum, where CGN-derived PFs synapse onto PCs.^19,21^

At PND6, we observed no genotype-dependent change in Vglut1 puncta coverage within the PC and molecular layers in *Scn2a^Y84X/+^* or *Scn2a^R1627X/+^*mice (Supplemental Fig. 5A-D; Supplemental Table 7), though there was a main effect of sex in mice from the R1627X line (LMEM: sex, F(D_1_,D_10.07_) = 5.34, *p* = 0.04). However, by PND12, *Scn2a^Y84X/+^* mice demonstrated a significant decrease in Vglut1 puncta coverage compared to wildtype littermates (LMEM: wildtype v. Y84X, genotype, F(D_1_,D_12.46_) = 6.89, *p* = 0.02). This was especially prevalent in cerebellar lobules IV/V and VII (LMEM: wildtype v. Y84X, genotype*lobule, F(D_7_,D_82.44_) = 2.21, *p* = 0.04; EMM: lobule IV/V, *t*(72.4) = 3.33, *p*□= 0.001; EMM: lobule VII, *t*(73.7) = 3.38, *p*□= 0.001; Fig. 3A,B; Supplemental Table 7). Conversely, *Scn2a^R1627X/+^* displayed no main effect of genotype or sex, and male *Scn2a^R1627X/+^* mice actually displayed an increase in Vglut1 puncta coverage compared to male wildtype in cerebellar lobules VIII and IX at PND12 (LMEM: wildtype v. R1627X, genotype*sex*lobule, F(D_7_,D_61.21_) =2.53, *p* = 0.02; EMM: lobule VIII, males, *t*(53.5) = −2.52, *p*□= 0.02; EMM: lobule IX, males, *t*(53.5) = −2.86, *p*□= 0.006; Fig. 3C,D; Supplemental Table 7). By PND21, no differences were observed in either *Scn2a^Y84X/+^* or *Scn2a^R1627X/+^*mice (Fig. 3E-H; Supplemental Table 7).

**Figure 3:**
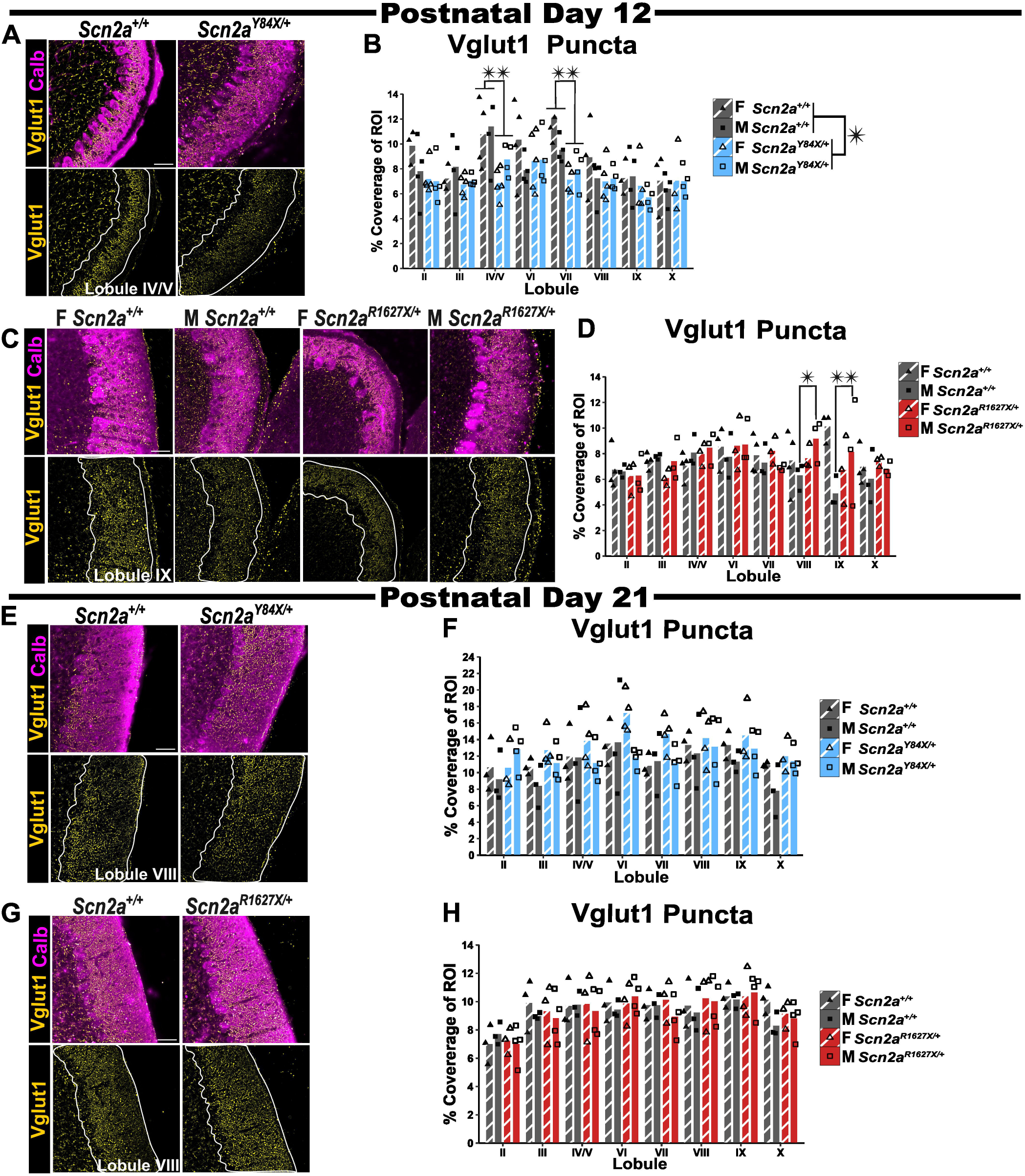
*Scn2a* PTC mice have variant- and age-specific differences in Vglut1 puncta distribution. **A)** Representative images of Vglut1 positive puncta (yellow) and Purkinje cells (magenta) from *Scn2a^Y84X/+^* mice and wildtype littermates at PND12. The region of interest (PC and molecular layer) is outlined in white. **B)** *Scn2a^Y84X/+^* mice have decreased Vglut1 puncta compared to wildtype (LMEM: wildtype v. Y84X, genotype, F(D_1_,D_12.46_) = 6.89, *p* = 0.02. This difference is most prominent in cerebellar lobules VI/V and VII (LMEM: wildtype v. Y84X, genotype*lobule, F(D_7_,D82.44) = 2.21, *p* = 0.04; EMM: lobule IV/V, t(72.40) = 3.33, *p* = 0.001; EMM: lobule VII, t(73.70) = 3.38, *p* = 0.001). **C)** Representative images of Vglut1 positive puncta (yellow) and Purkinje cells (magenta) from *Scn2a^R1627X/+^* mice and wildtype littermates at PND12. The region of interest (PC and molecular layer) is outlined in white. **D)** Male *Scn2a^R1627X/+^* mice have increased Vglut1 puncta compared to wildtype in cerebellar lobules in VIII and IX (LMEM: wildtype v. R1627X, genotype*sex*lobule, F(D_7_,D_61.21_ = 2.53, *p* = 0.02; EMM: Lobule VIII males, R1627X, t(53.50) = −2.52, *p* = 0.02; EMM: lobule IX, males, t(53.50) = −2.86, *p* = 0.006). **E)** Representative images of Vglut1 positive puncta (yellow) and Purkinje cells (magenta) from *Scn2a^Y84X/+^* mice and wildtype littermates at PND21. The region of interest (PC and molecular layer) is outlined in white. **F)** *Scn2a^Y84X/+^* mice have a similar Vglut1 expression pattern compared to wildtype littermates at PND21 (LMEM: wildtype v. Y84X, genotype, F(D_1_,D_10.08_) = 2.70, *p* = 0.13). **G)** Representative images of Vglut1 positive puncta (yellow) and Purkinje cells (magenta) from *Scn2a^R1627X/+^* mice and wildtype littermates at PND21. The region of interest (PC and molecular layer) is outlined in white. **F)** *Scn2a^R1627X/+^* mice had a similar Vglut1 expression pattern compared to wildtype littermates at PND21 (LMEM: wildtype v. R1627X, genotype, F(D_1_,D_5.57_) = 0.003, *p* = 0.96). Representative images were taken at 40X magnification. The scale bars are 50μm. Data are represented as mean +/- SEM. N = 3-5 per sex/lobule/genotype. (PTC = premature termination codon, LMEM = linear mixed effects model, EMM = estimated marginal means, PND = postnatal day)

As the cerebellar circuit matures, climbing fibers (CF), which arise from the inferior olive and synapse on PCs, undergo collateral pruning to establish a 1-to-few CF-PC relationship.^35,36^ Late CF pruning (≥PND12) is regulated by PF-PC activity.^36,37^ Therefore, we assessed Vglut2 expression, a CF marker at PND21, to determine whether *Scn2a* PTC mice retain extranumerary CF synapses. *Scn2a^Y84X/+^*mice had greater Vglut2 puncta coverage within the cerebellar molecular and PC layers compared to wildtype (LMEM: wildtype v. Y84X, genotype, F(D_1_,D_11.04_) = 19.24, *p* = 0.001; Fig. 4A,B; Supplemental Table 8). Conversely, *Scn2a^R1627X/+^* mice had no change in Vglut2 expression pattern compared to wildtype littermates (Fig. 4C,D; Supplemental Table 8).

**Figure 4:**
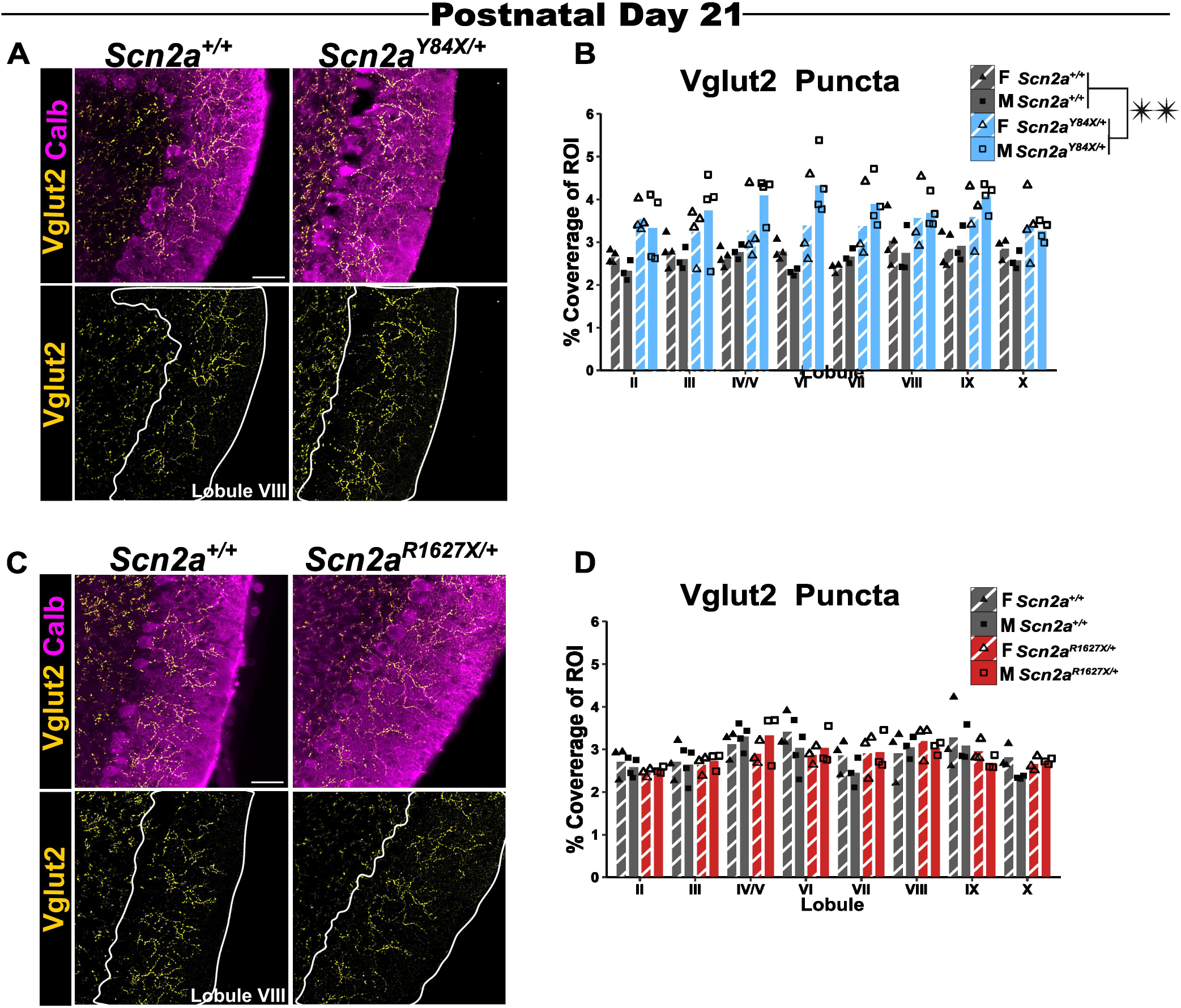
*Scn2a* mutant PTC mice have variant-specific changes in Vglut2 expression. **A)** Representative images of Vglut2 positive puncta (yellow) and Purkinje cells (magenta) from *Scn2a^Y84X/+^* mice and wildtype littermates at PND21. The region of interest (PC and molecular layer) is outlined in white. **B)** *Scn2a^Y84X/+^* mice have increased Vglut2 puncta compared to wildtype littermates at PND21 (LMEM: wildtype v. Y84X, genotype, F(D_1_,D_11.04_) = 19.24, *p* = 0.001). **C)** Representative images of Vglut2 positive puncta from *Scn2a^R1627X/+^* mice and wildtype littermates at PND21. The region of interest (PC and molecular layer) is outlined in white. **D)** *Scn2a^R1627X/+^* mice have similar Vglut2 puncta expression compared to wildtype littermates (LMEM: wildtype v. R1627X, genotype, F(D_1_,D_8.77_ = 0.20, *p* = 0.67). Representative images were taken at 40X magnification. The scale bar is 50μm. Data are represented as mean +/- SEM. N = 3-4 per sex/lobule/genotype. (PTC = premature termination codon, LMEM = linear mixed effects model, EMM = estimated marginal means, PND = postnatal day)

Since our histological analyses focused on the vermis, we also examined total levels of cerebellar presynaptic excitatory markers via capillary immunoassay. We detected reduced cerebellar Vglut1 protein expression in *Scn2a^Y84X/+^* mice at PND21 (LMEM: wildtype v. Y84X, genotype, F(D_1_,D_7.58_) = 5.71, *p* = 0.05), but no difference was seen in *Scn2a^R1627X/+^*mice (Fig. 5A-D; Supplemental Table 9). *Scn2a^Y84X/+^* mice also had increased cerebellar Vglut2 protein expression compared to wildtype (LMEM: wildtype v. Y84X, genotype, F(D_1_,D_4.77_) = 14.05, *p* = 0.02); this increase was prominently seen in male *Scn2a^Y84X/+^* mice compared to male wildtype (LMEM: wildtype v. Y84X, genotype*sex, F(D_1_,D_5.04_) = 16.92, *p* = 0.01; EMM: males, t(5.81) = −5.21, *p* = 0.002; Fig. 5E,F; Supplemental Table 9). Female mice in the Y84X line also had greater Vglut2 expression compared to male littermates (LMEM: sex, F(D_1_,D_4.61_) = 25.21, *p* = 0.01). In contrast, *Scn2a^R1627X/+^* mice had decreased cerebellar Vglut2 protein expression (LMEM: wildtype v. R1627X, genotype, F(D_1_,D_7.91_) = 7.20, *p* = 0.03; Fig. 5G,H; Supplemental Table 9).

**Figure 5:**
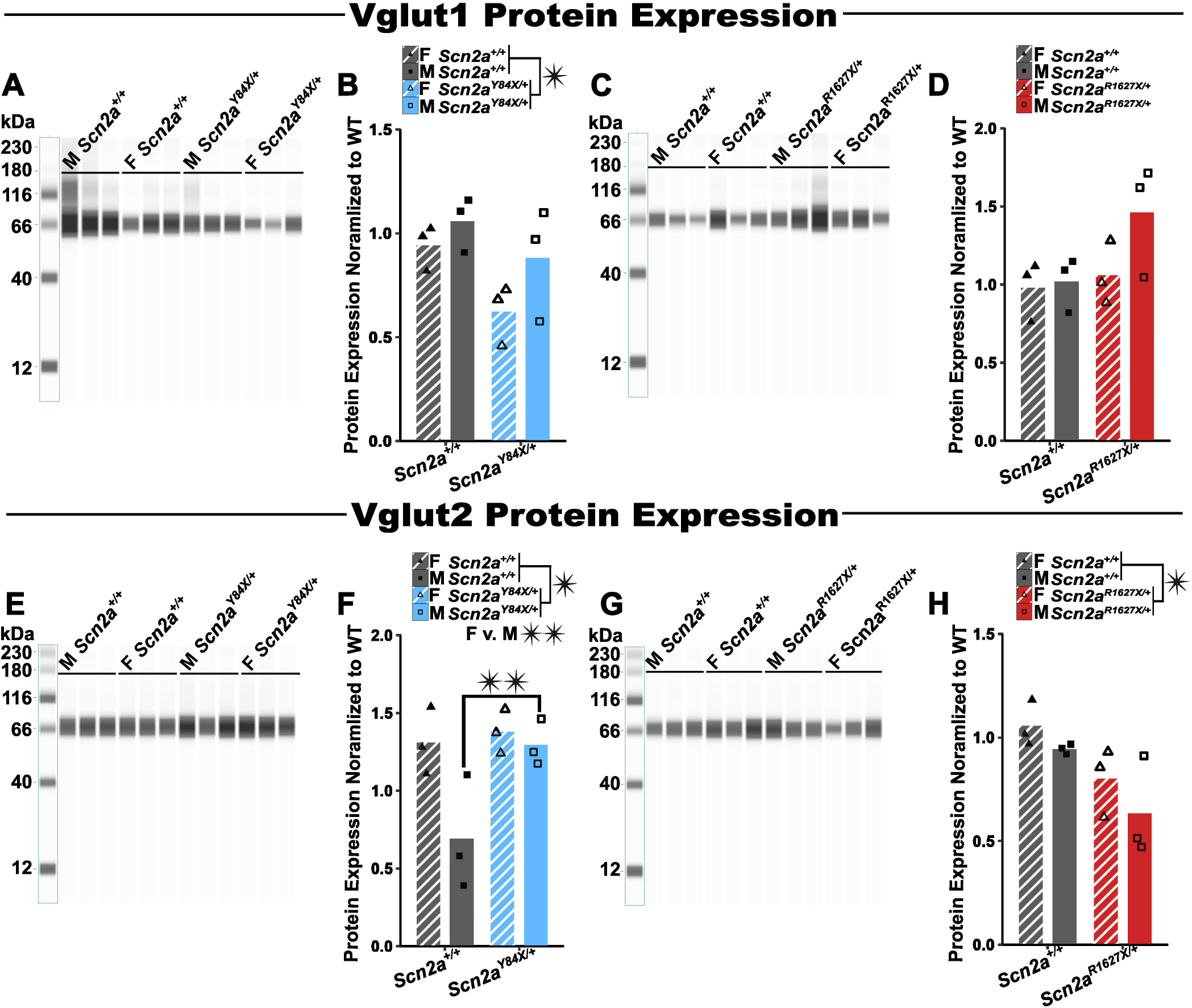
*Scn2a* mutant PTC mice have variant-specific alterations in Vglut1 and Vglut2 protein expression. **A)** Representative capillary immunoblot of cerebellar Vglut1 protein expression from *Scn2a^Y84X/+^* mice and wildtype littermates at PND21. **B)** *Scn2a^Y84X/+^* mice express less cerebellar Vglut1 protein than wildtype littermates (LMEM: wildtype v. Y84X, genotype, F(D_1_,D_7.58_) = 5.71, *p* = 0.05). **C)** Representative capillary immunoblot of cerebellar Vglut1 protein expression from *Scn2a^R1627X/+^* mice and wildtype littermates at PND21. **D)** *Scn2a^R1627X/+^* mice have similar Vglut1 protein expression compared to wildtype littermates at PND21 (LMEM: wildtypev. R1627X, genotype, F(D_1_,D_7.98_) = 1.66, *p* = 0.23). **E)** Representative JESS automated western blot of cerebellar Vglut2 protein expression from *Scn2a^Y84X/+^* mice and wildtype littermates at PND21. **F)** *Scn2a^Y84X/+^* mice have higher expression of Vglut2 protein in the cerebellum at PND21 compared to wildtype littermates (LMEM: wildtype v. Y84X, genotype, F(D_1_,D_4.77_) = 14.05, *p* = 0.02. This is most apparent between Male *Scn2a^Y84X/+^* mice and male wildtype littermates (LMEM: wildtype v. Y84X, genotype*sex, F(D_1_,D_5.04_) = 16.92, *p* = 0.01, EMM: males, t(5.81) = −5.21, *p* = 0.002). Female mice, in general also had increased Vglut2 cerebellar protein expression compared to male mice (LMEM: sex, F(D_1_,D_4.61_) = 25.21, *p* = 0.01; EMM: female v. male, t(5.36) = 4.87, *p* = 0.004). **G)** Representative JESS automated western blot of cerebellar Vglut2 protein expression from *Scn2a^R1627X/+^* mice and wildtype littermates at PND21. **H)** Evaluation of Vglut2 cerebellar protein expression in *Scn2a^R1627X/+^* mice revealed a decrease in Vglut2 cerebellar protein compared to wildtype littermates (LMEM: wildtype v. R1627X, genotype, F(D_1_,D_7.91_) = 7.20, *p* = 0.03). Data are represented as mean +/- SEM. Protein expression was normalized to wildtype. N = 3 per sex/genotype. (PTC = premature termination codon, LMEM = linear mixed effects model, EMM = estimated marginal means, PND = postnatal day).

### *Scn2a* PTCs differentially alter Purkinje cell density and soma size

Since CGNs express Na_V_1.2 and are a major excitatory input to PCs, we investigated whether *Scn2a* PTCs alter PC morphology during early-postnatal development. Given the dependence of PC maturation and morphology on CGN activity,^21,22^ we hypothesized that *Scn2a* PTC mice would have reduced PC soma area and density. At PND12, *Scn2a^Y84X/+^*mice had significantly lower PC density compared to wildtype (LMEM: wildtype v. Y84X, genotype, F(D_1_,D_11.16_) = 5.75, *p* = 0.04; Fig. 6A,B; Supplemental Table 10). On the other hand, *Scn2a^R1627X/+^* mice displayed largely unchanged PC density, with the exception of an increase in PC density between female *Scn2a^R1627X/+^*mice and female wildtype littermates in cerebellar lobule III at PND12 (LMEM: wildtype v. R1627X, genotype*sex*lobule, F(D_7_,D_55.12_) = 2.16, *p* = 0.05; EMM: lobule III, females, *t*(42.5) = −2.61, *p*□= 0.01; Fig. 6C,D; Supplemental Table 10). Additionally, at PND12, *Scn2a^Y84X/+^* mice had smaller PC somas compared to wildtype (LMEM: wildtype v. Y84X, genotype, F(D_1_,D_7.25_) =10.72, *p* = 0.01; Fig. 6E,F; Supplemental Table 11).

**Figure 6:**
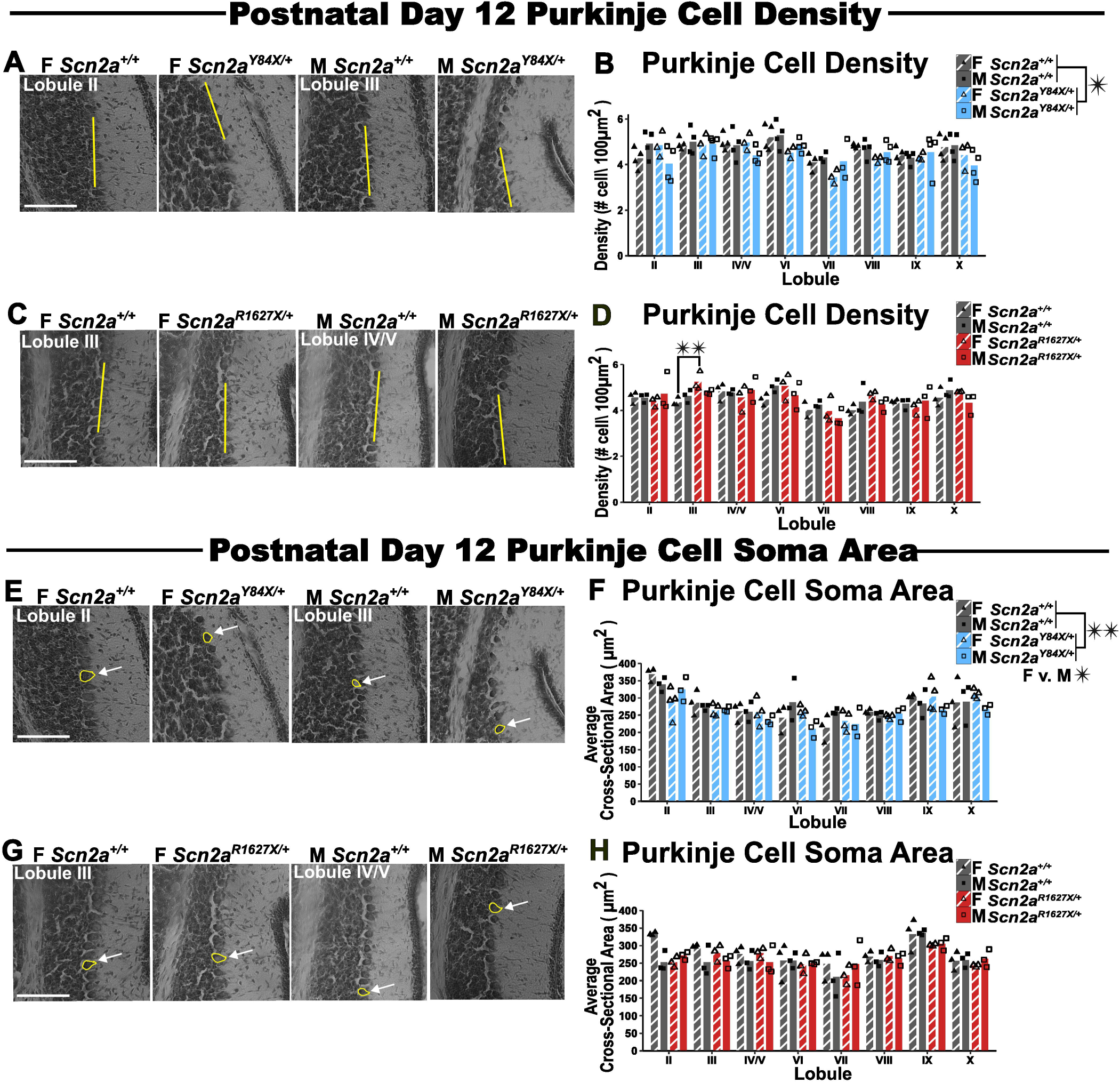
*Scn2a* PTC mutant mice exhibit variant specific changes in Purkinje cell morphology compared to wildtype littermates at PND12. **A)** Representative images of Purkinje cell density measurements from *Scn2a^Y84X/+^* mice and wildtype littermates at PND12. The yellow line represents the straight-line tool in imageJ used to take the measurement. B) *Scn2a^Y84X/+^* mice have decreased Purkinje cell density compared to wildtype littermates at PND12 (LMEM: wildtype v. Y84X, genotype, F(D_1_,D_11.16_) = 5.57, *p* = 0.04. C) Representative images of Purkinje cell density measurements from *Scn2a^R1627X/+^* mice and wildtype littermates at PND12. The yellow line represents the straight-line tool in imageJ used to take the measurement. D) Female *Scn2a^R1627X/+^* mice demonstrated increased Purkinje cell density in cerebellar lobule III compared to female wildtype mice at PND12 (LMEM: wildtype v. R1627X, genotype*sex*lobule, F(D_7_,D_55.12_) = 2.16, *p* = 0.05; EMM: lobule III females, t(42.50) = −2.61, *p* = 0.01). **E)** Representative images from *Scn2a^Y84X/+^* mice and wildtype littermates at PND12 of the cross-sectional area of Purkinje cell soma measurements. The white arrow is identifying how the polygon tool in imageJ was used to take the measurement. F) *Scn2a^Y84X/+^* mice had smaller Purkinje cell soma size compared to wildtype littermates at PND12 (LMEM: wildtype v. Y84X, genotype, F(D_1_,D_7.25_) = 10.72, *p* = 0.01. Female mice, in general, were found to have larger Purkinje cell somas than male littermates (LMEM: sex, F(D_1_,D_9.91_) = 5.21, *p* = 0.05). **G)** Representative images from *Scn2a^R1627X/+^* mice and wildtype littermates at PND12 of the cross-sectional area of Purkinje cell soma measurements. The white arrow is pointing at how the polygon tool in imageJ was used to take the measurement. H) *Scn2a^R1627X/+^* mice had similar Purkinje cell soma size compared to wildtype littermates at PND12 (LMEM: wildtype v. R1627X, genotype, F(D_1_,D_7.20_) = 0.22, *p* = 0.65). Representative images were taken at 20X magnification. The scale bar is 100μm. Data are represented as mean +/- SEM. N = 3-4 per sex/lobule/genotype. (PTC = premature termination codon, LMEM = linear mixed effects model, EMM = estimated marginal means, PND = postnatal day, PC = Purkinje cell)

Female mice in the Y84X line also possessed larger PCs than male mice (LMEM: sex, F(D_1_,D_9.91)_ = 5.21, *p* = 0.05). Consistent with the PC density data, there was no difference in PC soma size between *Scn2a^R1627X/+^* animals and wildtype at PND12 (Fig. 6G,H; Supplemental Table 11).

At PND21, both *Scn2a^Y84X/+^* and *Scn2a^R1627X/+^* mice demonstrated reduced PC density compared to respective wildtype littermates (LMEM: wildtype v. Y84X, genotype, F(D_1_,D_72_) = 21.27, *p* = <0.001; LMEM: wildtype v. R1627X, genotype, F(D_1_,D_6.69_) = 10.53, *p* = 0.02; Fig. 7A-D; Supplemental Table 10). *Scn2a^Y84X/+^* mice also continued to have smaller PC somas compared to wildtype littermates at PND21 (LMEM: wildtype v. Y84X, genotype, F(D_1_,D_8.30_) = 6.96, *p* = 0.03). This was most evident in cerebellar lobule II, where female *Scn2a^Y84X/+^* mice had smaller PC somas, and lobule VI, where male *Scn2a^Y84X/+^* mice had smaller PC somas compared to wildtype (LMEM: wildtype v. Y84X, genotype*sex*lobule, F(D_7_,D_63_) = 2.82, *p* = 0.01; EMM: lobule II, females, *t*(42.2) = 3.93, *p*□= 0.0003; EMM: lobule VI, males, *t*(51.7) = 3.51, *p*□= 0.0009; Fig. 7E,F; Supplemental Table 11). Interestingly, we observed that by PND21, *Scn2a^R1627X/+^* mice also exhibited decreased PC soma size compared to wildtype as well as an overall sex difference (LMEM: wildtype v. R1627X, genotype, F(D_1_,D_6.34_) = 6.59, *p* = 0.04; LMEM: sex, F(D_1_,D _6.62_) = 5.94, *p* = 0.05; Fig. 7G,H; Supplemental Table 11).

**Figure 7:**
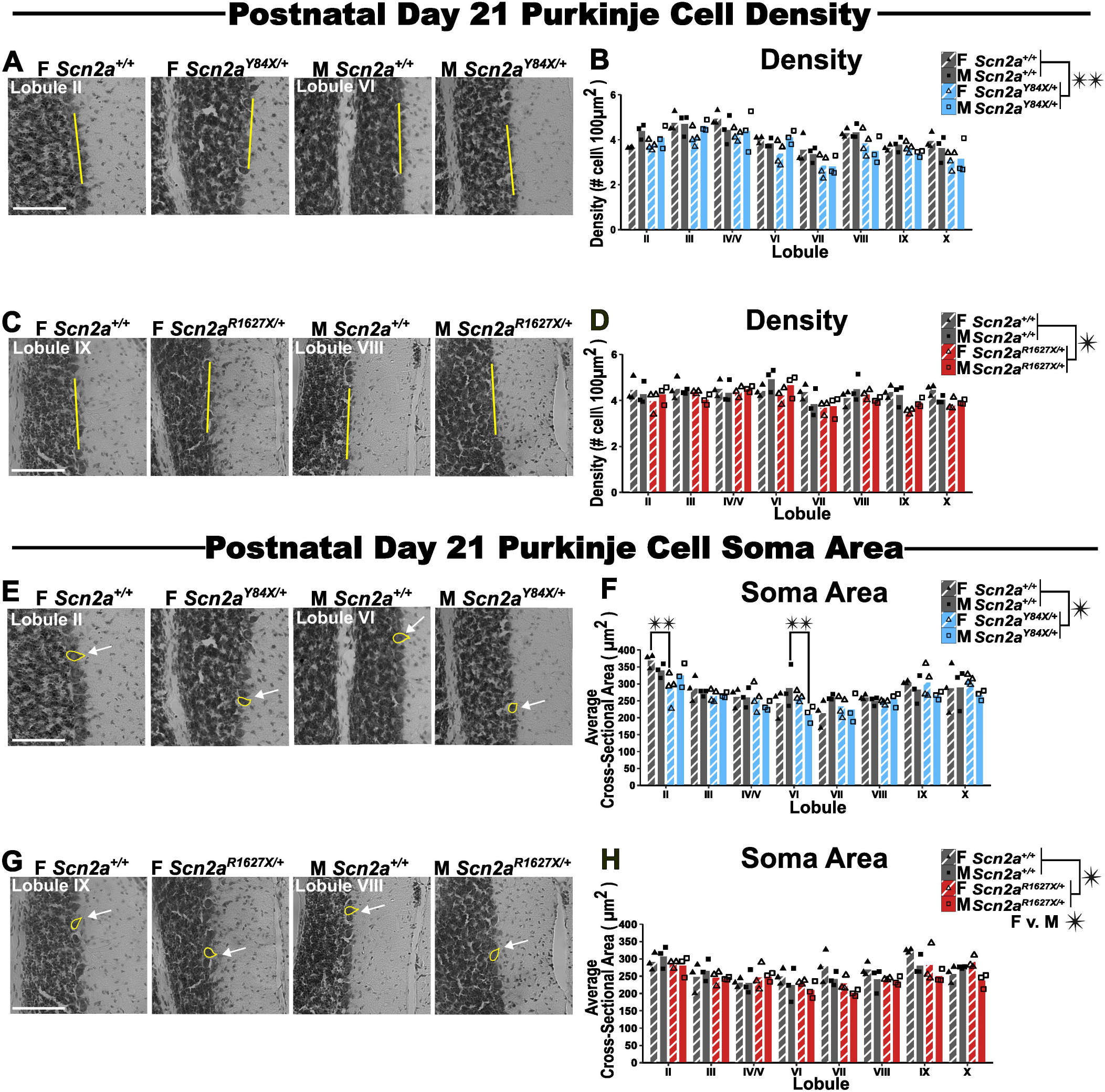
*Scn2a* PTC mutant mice exhibit variant specific changes in Purkinje cell morphology compared to wildtype littermates at PND21. **A)** Representative images of the 2-dimensional Purkinje cell density measurements from *Scn2a^Y84X/+^* mice and wildtype littermates at PND21. The yellow line represents the straight-line tool in imageJ used to take the measurement. B) *Scn2a^Y84X/+^* mice had decreased Purkinje cell density compared to wildtype littermates at PND21 (LMEM: wildtype v. Y84X, genotype, F(D_1_,D_72_) = 21.27, *p* < 0.001. **C)** Representative images of the 2-dimensional Purkinje cell density measurements from *Scn2a^R1627X/+^* mice and wildtype littermates at PND21. The yellow line represents the straight-line tool in imageJ used to take the measurement. D) *Scn2a^R1627X/+^* mice had decreased Purkinje cell density compared to wildtype littermates at PND21 (LMEM: wildtype v. R1627X, genotype, F(D_1_,D_6.69_) = 10.53, *p* = 0.02. **E)** Representative images from *Scn2a^Y84X/+^* mice and wildtype littermates at PND21 of the cross-sectional area of Purkinje cell soma measurements. The white arrow is identifying how the polygon tool in imageJ was used to take the measurement. F) *Scn2a^Y84X/+^* mice had smaller Purkinje cell somas than wildtype littermates (LMEM: wildtype v. Y84X, genotype, F(D_1_,D_8.30_) = 6.96, p = 0.03). This is most prominently seen in Female *Scn2a^Y84X/+^* mice compared to female wildtype littermates in cerebellar lobule II at PND21 (LMEM: wildtype v. Y84X, genotype*sex*lobule, F(D_7_,D_63_) = 2.82, p = 0.01; EMM: lobule II, females, t(42.20) = 3.93, *p* = 0.0003), and male Scn*2a^Y84X/+^* mice in cerebellar lobule VI compared to male wildtype littermates (EMM: lobule VI, males, t(57.70) = 3.51, *p* = 0.0009). **G)** Representative images from *Scn2a^R1627X/+^* mice and wildtype littermates at PND21 of the cross-sectional area of Purkinje cell soma measurements. The white arrow is pointing at how the polygon tool in imageJ was used to take the measurement. H) *Scn2a^R1627X/+^* demonstrated smaller Purkinje cell somas than wildtype littermates at PND21 (LMEM: wildtype v. R1627X, genotype, F(D_1_,D_6.34_) = 6.59, *p* = 0.04). Also, Female mice, in general, were found to have larger Purkinje cell somas than male littermates (LMEM: sex, F(D_1_,D_6.62_) = 5.94, p = 0.05). Representative images were taken at 20X magnification. The scale bar is 100μm. Data are represented as mean +/- SEM. N = 3-4 per sex/lobule/genotype. (PTC = premature termination codon, LMEM = linear mixed effects model, EMM = estimated marginal means, PND = postnatal day, PC = Purkinje cell)

### Developmental cerebellar layer dysmorphology in *Scn2a* PTC mice

Since CGN migration and maturation are activity-dependent,^21–23^ we assessed whether the external cerebellar granule layer (EGL) was altered in *Scn2a* PTC mice. The EGL is a postnatal structure which disappears as CGNs migrate to their terminal location in the internal granule layer. We hypothesized that *Scn2a* PTCs would impair CGN migration, resulting in prolonged retention of the EGL, manifesting as a thicker EGL at later ages. At PND6, we observed a genotype*sex interaction for EGL thickness in *Scn2a^Y84X/+^* mice (LMEM: wildtype v. Y84X, genotype*sex, F(D_1_,D_56.41_) = 4.79, *p* =0.03); however, follow up testing revealed no differences between sex and genotype groups (Fig. 8A,B; Supplemental Table 12). Male *Scn2a^R1627X/+^* mice had a thinner EGL compared to wildtype males in cerebellar lobule III, while female *Scn2a^R1627X/+^*mice had a thinner EGL compared to wildtype female mice in cerebellar lobule VIII at PND6 (LMEM: wildtype v. R1627X, genotype*sex*lobule, F(D_7_,D_73.33_) = 2.91, *p* = 0.01; EMM: lobule III, males, *t*(52.6) = 3.05, *p*□= 0.004; EMM: lobule VIII, females, *t*(57.8) = 2.86, *p*□= 0.01; Fig. 8C,D; Supplemental Table 12).

**Figure 8:**
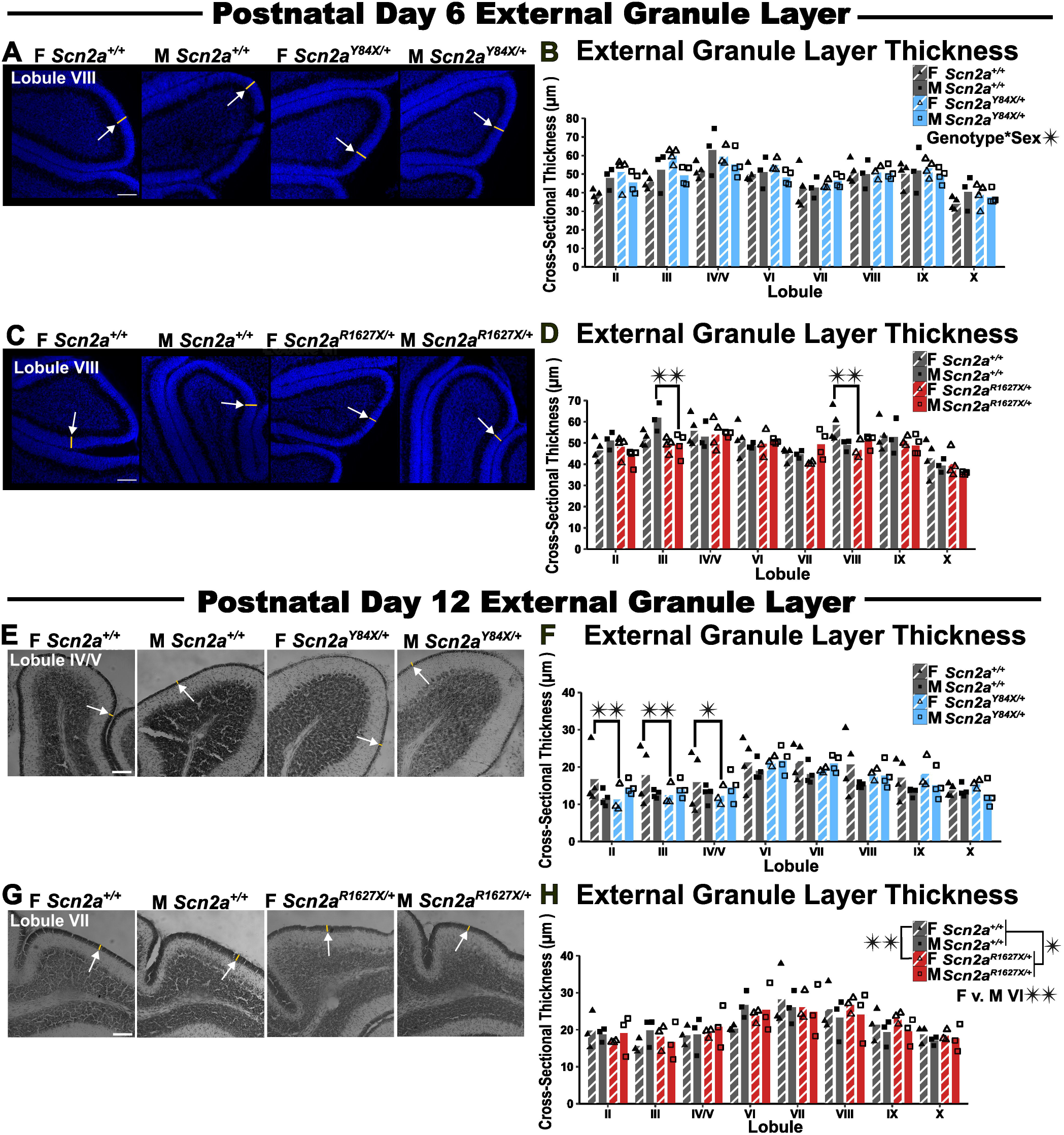
*Scn2a* PTC mutant mice demonstrate sex and age specific decreases in external granule layer thickness compared to wildtype littermates. **A)** Representative images of external granule layer thickness measurements from *Scn2a^Y84X/+^* mice and wildtype littermates at PND6. The white arrow is pointing to the straight-line tool in ImageJ used to take the measurement. **B)** Evaluation of PND6 *Scn2a^Y84X/+^* mice compared to wildtype littermates revealed an interaction effect between genotype and sex (LMEM: wildtype v. Y84X, genotype*sex, F(D_1_,D_56.41_) = 4.79, *p* = 0.03. Follow up testing revealed no differences (EMM: females, t(73.30) = −1.62, *p* = 0.11; EMM: males, t(65.04) = 1.50, *p* =0.14). **C)** Representative images of external granule layer thickness measurements from *Scn2a^R1627X/+^* mice and wildtype littermates at PND6. The white arrow is pointing at the straight-line tool in ImageJ used to take the measurement. **D)** Male *Scn2a^R1627X/+^* mice had a thinner external granule layer in lobule III and female *Scn2a^R1627X/+^* mice had a thinner external granule layer in lobule VIII compared to wildtype littermates at PND6. (LMEM: wildtype v. R1627X, genotype*sex*lobule, F(D_7_,D_73.33_) = 2.92, *p* = 0.01; EMM: lobule III males, t(52.60) = 3.05, *p* = 0.004; EMM: lobule VIII females, t(57.80) = 2.86, *p* = 0.01). **E)** Representative images from *Scn2a^Y84X/+^* mice and wildtype littermates at PND12 of the 2-dimensional external granule layer measurements. The white arrow is pointing at how the straight-line tool in imageJ was used to take the measurement. **F)** Female *Scn2a^Y84X/+^* mice had thinner external granule layers in cerebellar lobule II, III, IV/V compared to female wildtype mice at PND12 (LMEM: wildtype v. Y84X, genotype*sex*lobule, F(D_7_,D_77_) = 2.14, *p* = 0.05; EMM: lobule II, females, t(29.70) = 2.99, *p* = 0.0); EMM: lobule III, females, ΔM = 7.44, t(29.70) = 2.98, *p* = 0.01; EMM: lobule IV/V females, t(29.70) = 2.31, *p* = 0.03). **G)** Representative images from *Scn2a^R1627X/+^* mice and wildtype littermates at PND12 of the 2-dimensional external granule layer measurements. The white arrow is pointing at how the straight-line tool in imageJ was used to take the measurement. H) *Scn2a^R1627X/+^* mice had thinner external granule layers compared to wildtype littermates at PND12 (LMEM: wildtype v. R1627X, genotype, F(D_1_,D_7.03_) = 9.07, *p* = 0.02). This was especially apparent for female *Scn2a^R1627X/+^* mice compared to female wildtype littermates (LMEM: wildtype v. R1627X, genotype*sex, F(D_1_,D_7.03_) = 7.23, *p* = 0.03; EMM: females, t(7.05) = 3.75, *p* = 0.01). Also, female mice, in general, also had a thinner external granule layer in lobule VI compared to male littermates (LMEM: sex*lobule, F(D_7_,D_56_) = 2.21, *p* = 0.05; EMM: lobule IV, female v. male, t(61.60) = −2.52, *p* = 0.01). Representative images were taken at 20X magnification. The scale bar is 100μm. Data are represented as mean +/- SEM. N = 3-4 per sex/lobule/genotype. (PTC = premature termination codon, PTC = premature termination codon, LMEM = linear mixed effects model, EMM = estimated marginal means, PND = postnatal day, EGL = external granule layer)

At PND12, female *Scn2a^Y84X/+^* mice displayed a thinner EGL compared to female wildtype mice in cerebellar lobules II, III, and IV/V (LMEM: wildtype v. Y84X, genotype*sex*lobule, F(D_7_,D_77_) = 2.14, *p* = 0.05; EMM: lobule II, females, *t*(29.7) = 2.99, *p*□= 0.01; EMM: lobule III, females, *t*(29.7) = 2.98, *p*□= 0.01; EMM: lobule IV/V, females, *t*(29.7) = 2.31, *p*□= 0.03; Fig. 8E,F; Supplemental Table 12). No difference was identified between the male wildtype and male *Scn2a^Y84X/+^* mice at PND12 (Supplemental Table 12). *Scn2a^R1627X/+^* mice also had a thinner EGL compared to wildtype at PND12 (LMEM: wildtype v. R1627X, genotype, F(D_1_,D_7.03_) = 9.07, *p* = 0.02; Fig 8G,H; Supplemental table 12). This difference was most prominent between female *Scn2a^R1627X/+^* and female wildtype littermates (LMEM: wildtype v. R1627X, genotype*sex, F(D_1_,D_7.03_) = 7.23, *p* = 0.03; EMM: females, *t*(7.05) = 3.75, *p*□= 0.007). Independent of genotype, females had a thinner EGL in lobule VI compared to males (LMEM: sex*lobule, F(D_1_,D_7.03_) = 7.23, *p* = 0.03; EMM: lobule VI, female v. male, *t*(61.6) = - 2.52, *p*□= 0.01).

Since we had detected smaller PC somas in some ages of *Scn2a* PTC mice, we evaluated if these changes also caused *Scn2a* PTC mice to have a thinner cerebellar molecular layer (ML). However, neither *Scn2a^Y84X/+^* or *Scn2a^R1627X/+^* mice at PND12 nor PND21 displayed differences in the cerebellar ML thickness compared to respective wildtype littermates (Supplemental Fig. 4A-I; Supplemental Table 13).

## Discussion

Our findings substantiate the cerebellum as an early site of *SCN2A*-related pathophysiology and reveal variant-dependent disruptions in cerebellar circuit development and behavior. The more pronounced phenotypes in *Scn2a^Y84X/+^* mice, including altered glutamatergic synaptic markers and cerebellar-dependent behaviors, suggests that the position and molecular consequences of *SCN2A* PTCs may influence the timing and severity of cerebellar dysfunction. Our work defines a developmental window wherein cerebellar dysfunction emerges.

Collectively, we demonstrate that *Scn2a* PTC variants are associated with unique molecular cerebellar development disruptions and distinct cerebellum-related behavioral phenotypes.

Behaviorally, while overall pup growth and non-cerebellar behaviors were largely preserved, *Scn2a^Y84X/+^*mice were faster to perform CA and SRR than wildtype littermates. This was most evident in male *Scn2a^Y84X/+^* mice who performed CA and SRR with shorter latencies, and demonstrated greater aptitude on SRR, than male wildtype counterparts. *Scn2a^R1627X/+^*mice did not demonstrate any cerebellar-related behavioral changes. These data illustrate differential behavioral impacts of distinct *Scn2a* PTCs.

Relatedly, we also observed sparser labeling of Vglut1 puncta in *Scn2a^Y84X/+^* mice at PND12, indicating fewer contacts from PFs onto PCs (Vglut1 is enriched where PFs synapse onto PC dendrites).^19,38^ This suggests that heterozygous Na_v_1.2 loss impairs the ability of PFs synapse formation. However, male *Scn2a^R1627X/+^* mice showed modest increases in Vglut1 puncta abundance, despite presumably having similar heterozygous cerebellar Na_v_1.2 loss. Our previous work evaluating cortical glutamatergic neurons may explain this disparity. We have shown that cortical glutamatergic neurons from *Scn2a^Y84X/+^* mice have more severe functional deficits than *Scn2a^R1627X/+^* neurons: *Scn2a^Y84X/+^*neurons display greater action potential velocity attenuation and a depolarized spike threshold than both *Scn2a^R1627X/+^* and wildtype neurons.^25^ If similar functional deficits occur in CGNs this could explain why we see presynaptic Vglut1 reductions in *Scn2a^Y84X/+^* mice and not *Scn2a^R1627X/+^*mice. Answering this hypothesis requires electrophysiological measurements in CGNs. Although neither *Scn2a^Y84X/+^* or *Scn2a^R1627X/+^* mice demonstrated a change in Vglut1 puncta distribution at PND21, *Scn2a^Y84X/+^* mice did express less cerebellar Vglut1 protein This indicates that our histological method for measuring excitatory synapse distribution may underestimate differences, since we can only reliably perform these measurements at the vermis, and excitatory synapses are distributed throughout the hemispheres as well. Overall, our data provide evidence for reduced Vglut1 expression in *Scn2a^Y84X/+^* mice but not *Scn2a^R1627X/+^* mice.

We additionally observed increased Vglut2 puncta in *Scn2a^Y84X/+^* mice, and increased total cerebellar Vglut2 protein in male *Scn2a^Y84X/+^* mice, at PND21. Late CF pruning (≥PND12) is heavily regulated by PF-PC activity, which our Vglut1 data suggest is reduced, while early CF pruning is driven by cell-intrinsic properties.^35,37,39^ As CFs are the predominant source of cerebellar Vglut2 after PND11,^38,40^ our results suggest delayed late CF pruning, which may stall the progression toward the expected 1-to-few relationship between CFs and PCs.^35^ Importantly, during the first two postnatal weeks, Vglut2 is the predominant presynaptic glutamatergic marker in the developing cerebellum, expressed by both PFs and CFs.^41^ Around PND11, Vglut1 is upregulated in PFs, with spatial distribution spanning from the proximal to distal edge of the molecular layer, while Vglut2 expression is relegated to CFs.^40,41^ Thus, another interpretation of these data is that PFs from *Scn2a^Y84X/+^* mice are delayed in their transition from Vglut2 expression to Vglut1. While *Scn2a^Y84X/+^*mice demonstrate sparser Vglut1 puncta at PND12, the spatial distribution matches that of wildtype. Furthermore, Vglut2, although increased in PND21 *Scn2a^Y84X/+^* mice, is not diffusely scattered across the molecular layer but is linearly organized along putative CFs, like wildtype littermates. Together, this suggests that increased Vglut2 puncta density and protein expression is driven by delayed CF pruning and/or retention of extranumerary CFs. In contrast, *Scn2a^R1627X/+^* mice displayed no changes in Vglut2 puncta compared to wildtype littermates but did demonstrate a reduction in total cerebellar Vglut2 protein at PND21. These data suggest that *Scn2a^R1627X/+^* mice do not show delayed CF pruning but perhaps demonstrate a mild increase in pruning. Further exploration of cerebellar Vglut1 and Vglut2 expression including timepoints prior to PND21 (a limitation of our study) coupled with experiments mapping CF structure would clarify these results.

Differences in behavioral phenotypes between the two *Scn2a* PTC lines may be influenced by the molecular changes identified. The plausible retention of aberrant CF collaterals in *Scn2a^Y84X/+^* mice, as suggested by the increased spatial and protein expression of Vglut2, would likely increase CF-PC input. CFs produce complex spikes which briefly suppress PC activity and are related to long term depression (LTD).^42,43^ Additionally, a recent study found that high-frequency burst stimulation of PFs resulted in reduced excitatory postsynaptic current amplitude in *Scn2a^+/-^* mice, indicating a diminution in PF-PC synaptic transmission and reduced excitatory input due to heterozygous Na_v_1.2 loss, impairing long-term potentiation (LTP).^22^ Collectively, heterozygous Na_v_1.2 loss leading to impaired LTP, reduced PF-PC contacts, and increased LTD, could lead to PC suppression and augment disinhibition of deep cerebellar nuclei, potentially explaining the speedier performance of *Scn2a^Y84X/+^* mice in cerebellar-related reflex behaviors.^42,44,45^ *Scn2a^R1627X/+^* mice appear quite different: reduced Vglut2 in *Scn2a^R1627X/+^*mice may correlate with reduced LTD, and although they may have a similar LTP impairment as *Scn2a^+/-^* mice, it is not coupled to reduced Vglut1, suggesting that the deep cerebellar nuclei of *Scn2a^R1627X/+^* mice are unlikely to be disinhibited. Our lack of cerebellar electrophysiology data limits our interpretations of these data. Measurements of PC and CF activity, alongside analysis of deep cerebellar nuclei, could help clarify how *SCN2A* PTCs alter cerebellar output and related motor behaviors.

Both *Scn2a* PTC models display reduced PC soma size and density. This is consistent with other ASD-related models, including valproate exposure models and *Shank3*-KO mice, along with postmortem human ASD brains.^12,46–50^ *Scn2a^Y84X/+^* mice display reduced PC soma size and density at PND12 and PND21, while *Scn2a^R1627X/+^*mice display similar PC changes only at PND21. These changes may result from the altered distribution and protein expression of presynaptic glutamatergic excitatory markers in these mice. Since *Scn2a^Y84X/+^* and *Scn2a^R1627X/+^* mice display different molecular and behavioral phenotypes, their similar PC morphologies do not explain the differences. We also did not observe changes in molecular layer thickness, which often accompanies altered PC dendritic arbors. Future studies should further investigate PC complexity and electrophysiology to elucidate how PCs, which have not been shown to express *SCN2A*,^51–53^ are impacted by *Scn2a* PTCs. While altered PC soma size and density may affect cerebellar function, they do not closely correlate with the behavioral differences reported here.

Contrary to our hypothesis, we generally observed thinner EGLs in *Scn2a* PTC mice. This suggests accelerated CGN migration but could also reflect fewer CGN progenitors or impaired proliferation. While our study does not directly address this, impaired CGN progenitor expression and proliferation is unlikely in *Scn2a* PTC mice, as mice with CGN progenitor ablation or silencing of early-postnatal cerebellar glutamatergic transmission exhibit severe tremor and altered USVs, which neither *Scn2a* PTC model demonstrates.^22,54^ Moreover, developmental models with complete CGN progenitor ablation or silenced cerebellar glutamatergic transmission display significant increases in SRR latency, whereas *Scn2a^Y84X/+^* mice demonstrate the opposite. In contrast, when most CGN excitatory neurotransmission is inhibited through Ca_V_2.1 channel deletion, gross motor skills are preserved while cerebellar-related behaviors are impaired; this is more similar to our findings from *Scn2a^Y84X/+^* mice.^55^

CGN migration, maturation, and signaling are integral to the development of the larger cerebellar cortex, therefore, early CGN migration could have expedited maturation, influencing some of the earlier identified changes. However, more specific metrics to assess the functional maturity of the cerebellar circuit would be necessary to investigate this idea.^22,23,56^

Our study further supports distinct phenotypes associated with *SCN2A* PTCs. While the precise basis for the differential impact of unique PTCs is nebulous, it may be related to PTC position within the gene which determines transcript susceptibility to nonsense-mediated decay (NMD). *Scn2a-*p.Y84X occurs early in the C-terminal domain while Scn2a-p.R1627X occurs later in the 4th transmembrane region. We recently demonstrated greater NMD in Y84X-containing transcripts compared to R1627X-containing transcripts in the brain.^25^ Consistent with this, an updated computational model for NMD in disease suggests that greater NMD may exacerbate *SCN2A* PTC phenotypes.^25,57^ This difference could contribute to the more prominent phenotypes observed in *Scn2a^Y84X/+^*mice compared to *Scn2a^R1627X/+^* mice. Establishing a developmental timeline and understanding of cerebellar impairment may inform earlier diagnosis of *SCN2A* LOF variants and coordinate timing of therapeutic intervention.

## Supporting information

Supplemental Stat Summary Tables

## Data availability

Data is available upon request to the corresponding author.

## Acknowledgements

A huge thanks goes to Madison O’Bryan for her assistance with mouse colony management and training RG on thionin staining. This work was inspired by our conversations with the FamilieSCN2A Foundation, Leah Schust, and Emily Park; a big thank you goes out to them for encouraging our efforts and being willing to answer our questions about the lived experience and concerns caretakers have about their loved ones with *SCN2A*-related disorders.

