## Supplementary Figures and Legends for "Developmental Cerebellar Pathology in Mouse Models of *SCN2A* Premature Termination Codon Variants"

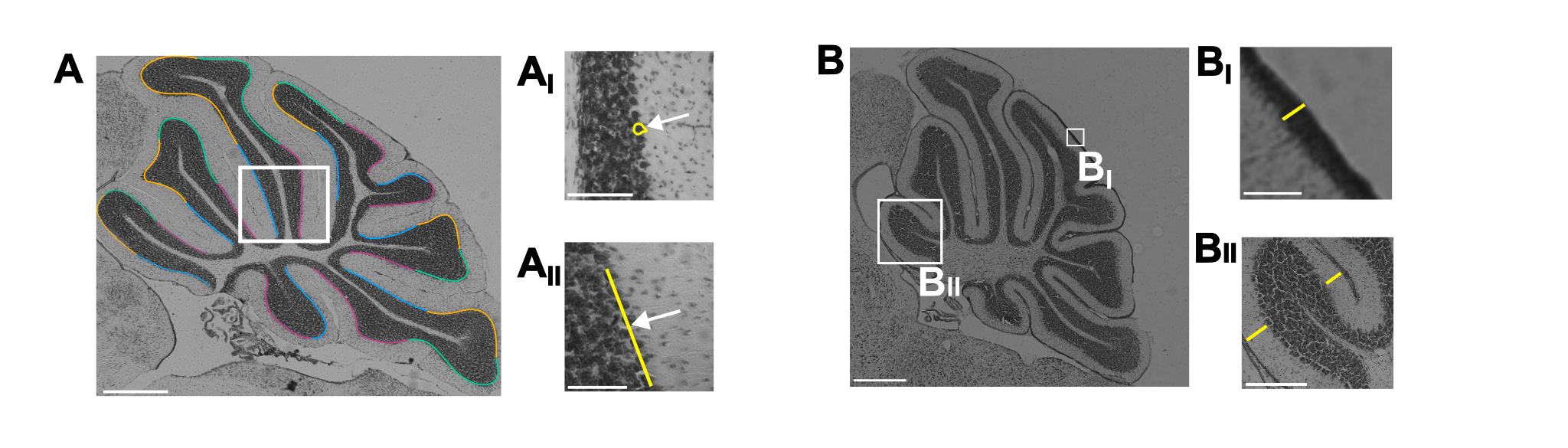

**Supplemental Figure 1: Representative images of how measurements were taken. A)** The colored lines represent the 2 to 4 regions that were used to take measurements from for histological analysis (Blue = anterior base, Orange = anterior crown, Pink = posterior base, Green = posterior crown). These regions were used across all ages analyzed for histology. **A_I_**) Represents a zoomed in region of A (the white square) the white arrow is pointing to an example of how the Purkinje cell soma area was traced in a vermal cerebellar section. **A_II_)** Represents a zoomed in region of A (the white square) with the white arrow pointing to an example of how Purkinje cell density was measured using the straightline tool in imageJ. B) Is a representative thionin image from a PND12 mouse. **B_I_**) Demonstrates an example of how the 2-dimensional thickness of the external granule layer was measured using the straightline tool in imageJ. This same process was used for DAPI stained images at PND6. **B_II_)** Showcases an example of how the 2-dimensional thickness of the molecular layer was measured at PND12 and PND21 using the straightline tool in imageJ. (PND = postnatal day)

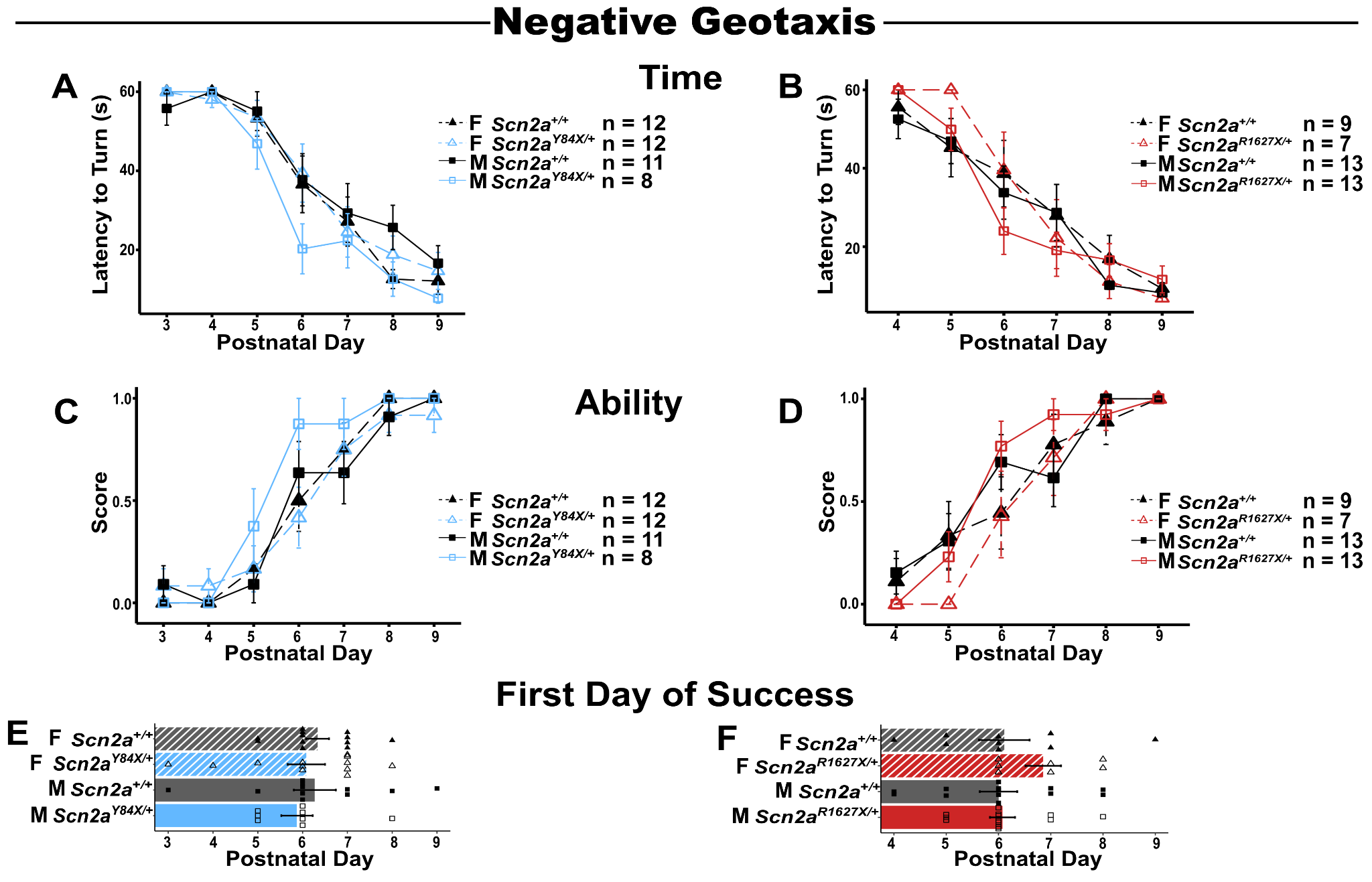

**Supplemental Figure 2: *Scn2a* mutant PTC mice have unaltered performance on negative geotaxis compared to wildtype littermates. A)** *Scn2a^Y84X/+^* mice and **B)** *Scn2a^R1627X/+^* mice perform the negative geotaxis reflex in a similar amount of time to their respective wildtype littermates (LMEM: wildtype v. Y84X, genotype, F(D_1_,D_36.13_) = 1.11, p = 0.30; LMEM: wildtype v. R1627X, genotype, F(D_1_,D_37.26_) = 1, *p* = 0.32). **C)** *Scn2a^Y84X/+^* mice and **D)** *Scn2a^R1627X/+^* mice achieve a similar score on the negative geotaxis reflex similar to their respective wildtype littermates (LMEM: wildtype v. Y84X, genotype, F(D_1_,D_36.19_) = 0.44, *p* = 0.51; LMEM: wildtype v. R1627X, genotype, F(D_1_,D_35.23_) = 0.002, p = 0.96). **E)** *Scn2a^Y84X/+^* mice and **F)** *Scn2a^R1627X/+^* mice first achieve the negative geotaxis reflex on a similar day to their respective wildtype littermates (LMEM: wildtype v. Y84X, genotype, F(D_1_,D_37.8_) = 0.16, *p* = 0.69; LMEM: wildtype v. R1627X, genotype, F(D_1_,D_35.61_) = 0.97, *p* = 0.33). Data are represented as mean +/- SEM. (PTC = premature termination codon, LMEM = linear mixed effects model, EMM = estimated marginal means)

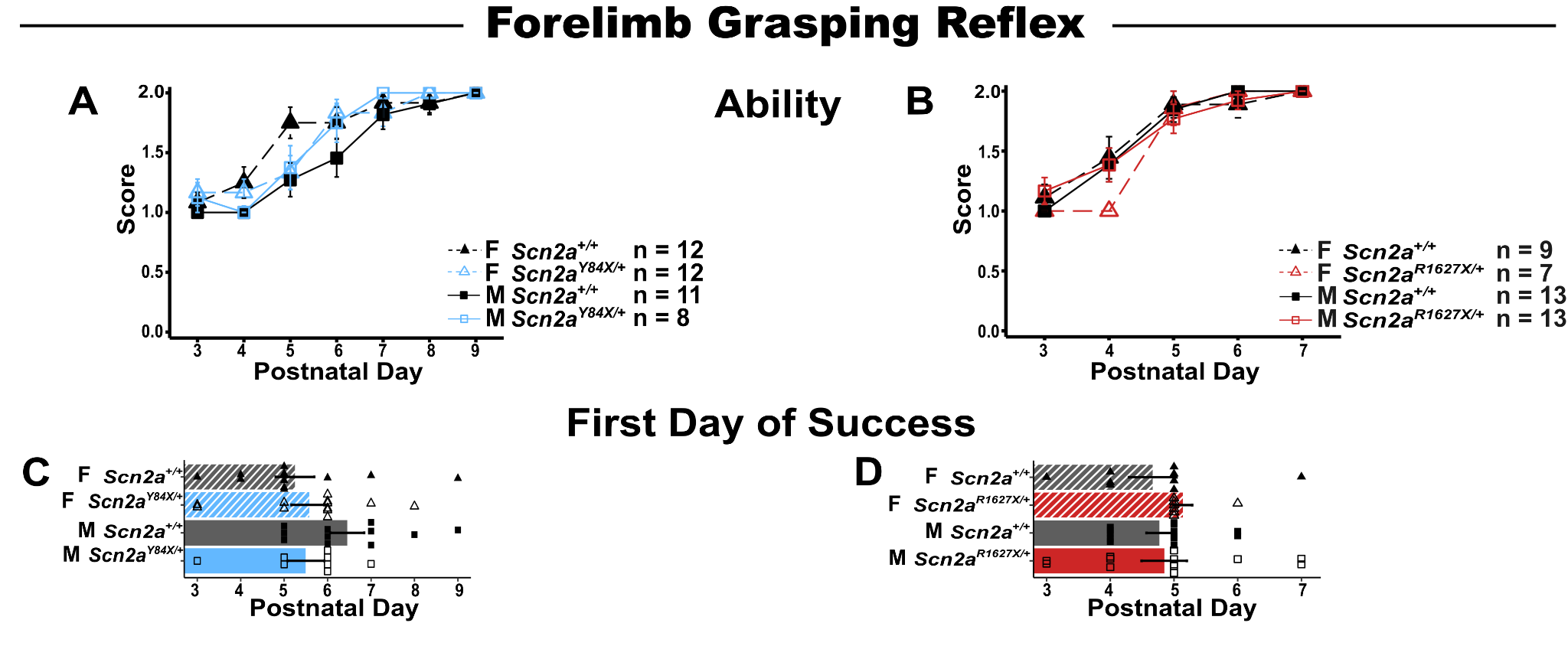

**Supplemental Figure 3: *Scn2a* mutant PTC mice have unaltered performance on the forelimb grasping reflex compared to wildtype littermates. A)** *Scn2a^Y84X/+^* mice and **B)** *Scn2a^R1627X/+^* mice perform the forelimb grasping reflex similar to their respective wildtype littermates (LMEM: wildtype v. Y84X, genotype, F(D_1_,D_34.86_) = 0.69, p = 0.41; LMEM: wildtype v. R1627X, genotype, F(D_1_,D_33.01_) = 0.003,, p= 0.96). **C)** *Scn2a^Y84X/+^* mice and **D)** *Scn2a^R1627X/+^* mice first achieve the forelimb grasping reflex on a similar day to their respective wildtype littermates (LMEM: wildtype v. Y84X, genotype, F(D_1_,D_38.98_) = 00.58, *p* = 0.45; LMEM: wildtype v. R1627X, genotype, F(D_1_,D_33.51_) = 00.02,*p* = 0.88). Data are represented as mean +/- SEM. (PTC = premature termination codon, LMEM = linear mixed effects model, EMM = estimated marginal means)

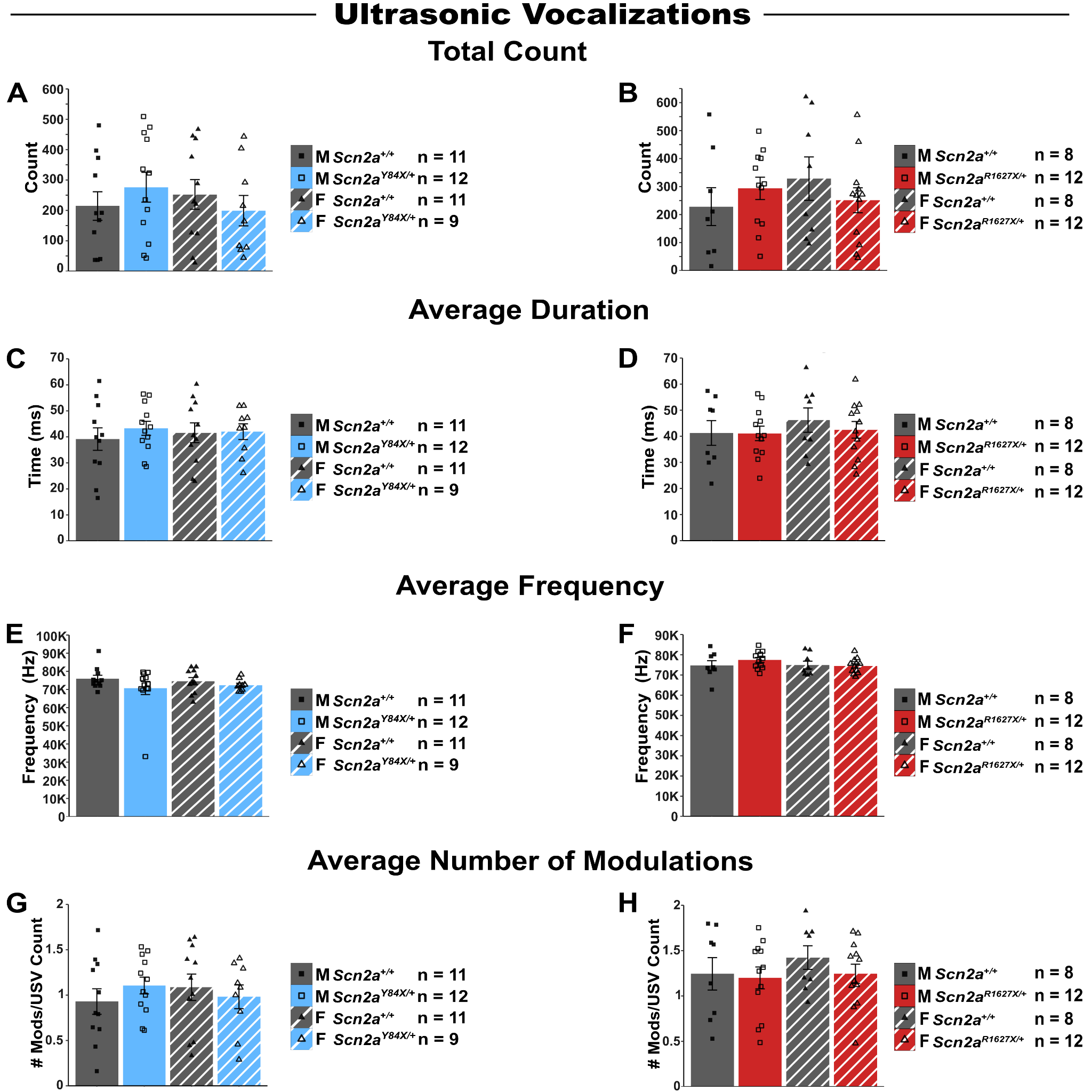

**Supplemental Figure 4: *Scn2a* mutant PTC mice have unaltered performance in ultrasonic vocalizations compared to wildtype littermates. A)** *Scn2a^Y84X/+^* mice and **B)** *Scn2a^R1627X/+^* mice produce a similar number of USVs as their respective wildtype littermates (LMEM: wildtype v. Y84X, genotype, F(D_1_,D_40.32_) = 0.02, p = 0.89; LMEM: wildtype v. R1627X, genotype, F(D_1_,D_33.30_) = 0.02, *p* = 0.89). **C)** *Scn2a^Y84X/+^* mice and **D)** *Scn2a^R1627X/+^* mice USVs are of a similar duration as their respective wildtype littermates (LMEM: wildtype v. Y84X, genotype, F(D_1_,D_40.99_) = 0.79, *p* = 0.38; LMEM: wildtype v. R1627X, genotype, F(D_1_,D_36_ ) = 0.27, *p* = 0.61 ). **E)** *Scn2a^Y84X/+^* mice and **F)** *Scn2a^R1627X/+^* mice USVs have a similar average frequency compared to their respective wildtype littermates (LMEM: wildtype v. Y84X, genotype, F(D_1_,D_42.44_) = 1.24, *p* = 0.27; LMEM: wildtype v. R1627X, genotype, F(D_1_,D_36_) = 0.49, *p* = 0.49). **G)** *Scn2a^Y84X/+^* mice and **H)** *Scn2a^R1627X/+^* mice produce an average number of modulations per USV similar to their respective wildtype littermates (LMEM: wildtype v. Y84X, genotype, F(D_1_,D_42.94_) = 0.770, *p* = 0.39; LMEM: wildtype v. R1627X, genotype, F(D_1_,D_34.31_) = 0.954, *p* = 0.34). Data are represented as mean +/- SEM. (PTC = premature termination codon, LMEM = linear mixed effects model, EMM = estimated marginal means, PND = postnatal day, USV = ultrasonic vocalization)

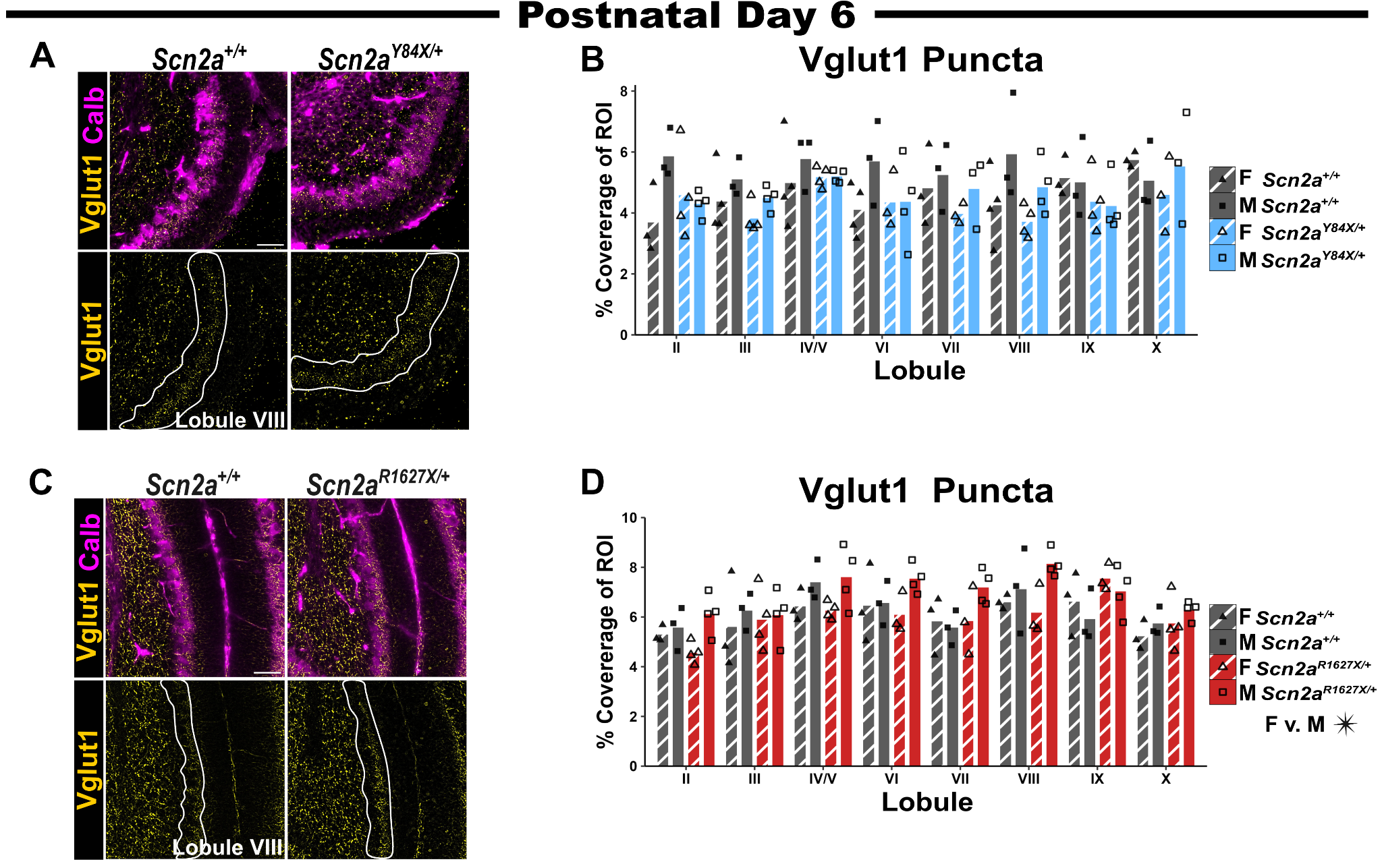

**Supplemental Figure 5: *Scn2a* mutant PTC mice have a similar Vglut1 puncta expression pattern to wildtype littermates at PND6. A)** Representative images from *Scn2a^Y84X/+^* mice and wildtype littermates of Vglut1 positive puncta (yellow) and Purkinje cells (magenta) at PND6. The region of interest (PC and molecular layer) is outlined in white. **B)** *Scn2a^Y84X/+^* mice had a similar expression pattern of Vglut1 positive puncta within the vermis as wildtype littermates (LMEM: wildtype v. Y84X, genotype, F(D_1_,D_10.78_) = 2.54, *p* = 0.14) **C)** Representative images from *Scn2a^R1627X/+^* mice and wildtype littermates of Vglut1 positive puncta (yellow) and Purkinje cells (magenta) at PND6. The region of interest (PC and molecular layer) is outlined in white. **D)** *Scn2a^R1627X/+^* mice had a similar expression pattern of Vglut1 positive puncta within the vermis as wildtype littermates (LMEM: wildtype v. R1627X, genotype, F(D_1_,D_10.07_) = 1.69, *p* = 0.22). However, male mice, in general, had increased Vglut1 positive puncta compared to female littermates (LMEM: sex, F(D_1_,D_10.07_) = 5.34, *p* = 0.04). Representative images were taken at 40X magnification. The scale bar is 50μm. Data are represented as mean +/- SEM. N = 3-4 per sex/lobule/genotype. (PTC = premature termination codon, LMEM = linear mixed effects model, EMM = estimated marginal means, PND = postnatal day)

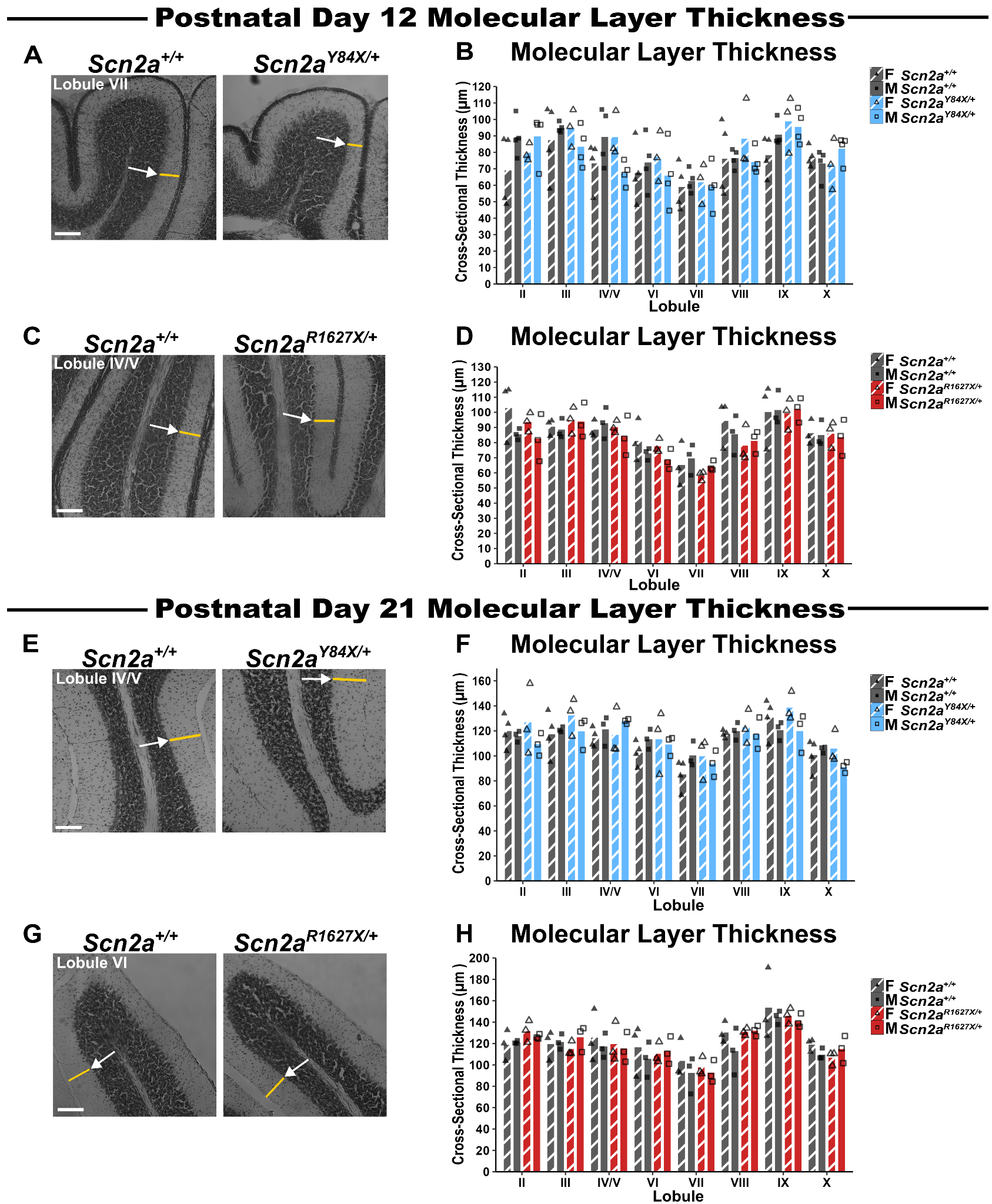

**Supplemental Figure 6: *Scn2a* mutant PTC mice have a similar 2-dimensional molecular layer thickness compared to wildtype littermates. A)** Representative images from *Scn2a^Y84X/+^* mice and wildtype littermates of PND12 of the 2-dimensional molecular layer measurements. The white arrow is pointing at how the straight-line tool in imageJ was used to take the measurement. **B)** *Scn2a^Y84X/+^* mice had a similar 2-dimensional molecular layer thickness as wildtype littermates at PND12 (LMEM: wildtype v. Y84X, genotype, F(D_1_,D_8.20_) = 0.60, *p* = 0.46). **C)** Representative images from *Scn2a^R1627X/+^* mice and wildtype littermates of PND12 of the 2-dimensional molecular layer measurements. **D)** *Scn2a^R1627X/+^* mice had a similar 2-dimensional molecular layer thickness as wildtype littermates at PND12 (LMEM: wildtype v. R1627X, genotype, F(D_1_,D_7.11_) = 0.03, *p* = 0.87). **E)** Representative images from *Scn2a^Y84X/+^* mice and wildtype littermates of PND21 of the 2-dimensional molecular layer measurements. The white arrow is pointing at how the straight-line tool in imageJ was used to take the measurement. **F)** *Scn2a^Y84X/+^* mice had a similar 2-dimensional molecular layer thickness as wildtype littermates at PND21 (LMEM: wildtype v. Y84X, genotype, F(D_1_,D_9_) = 0.39, *p* = 0.55). **G)** Representative images from *Scn2a^R1627X/+^* mice and wildtype littermates of PND21 of the 2-dimensional molecular layer measurements. **H)** *Scn2a^R1627X/+^* mice had a similar 2-dimensional molecular layer thickness as wildtype littermates at PND21 (LMEM: wildtype v. R1627X, genotype, F(D_1_,D_23.83_) = 0.02, *p* = 0.90). Representative images were taken at 20X magnification. The scale bar is 100μm. Data are represented as mean +/- SEM. (LMEM = linear mixed effects model, EMM = estimated marginal means, PND = postnatal day)
