## Supplemental Stat Summary Tables for "Developmental Cerebellar Pathology in Mouse Models of *SCN2A* Premature Termination Codon Variants"

**Supplemental Table 1. Effects of the Y84X and R1627X variants on early postnatal health measurements in mice**

| **Weight** | | | | | |
| --- | --- | --- | --- | --- | --- |
| Measure | Mouse Line | Effect | NumDF,DenDF | F-Value | *P*-Value |
| Weight | Y84X | Genotype | 1, 38.51 | 0.19 | 0.66 |
|  |  | Sex | 1, 34.9 | 0.04 | 0.84 |
|  |  | Day | 11, 429 | 536.63 | **<0.001** |
|  |  | Genotype* Sex | 1, 37.31 | 0.55 | 0.46 |
|  |  | Genotype* Day | 11, 429 | 1.42 | 0.16 |
|  |  | Sex* Day | 11, 429 | .33 | 0.98 |
|  |  | Genotype*Sex*Day | 11, 429 | .66 | 0.78 |
|  | R1627X | Genotype | 1, 33.69 | 1.94 | 0.17 |
|  |  | Sex | 1, 32.81 | 0.52 | 0.47 |
|  |  | Day | 11, 418 | 636 | **<0.001** |
|  |  | Genotype* Sex | 1, 33.52 | 3.07 | 0.09 |
|  |  | Genotype* Day | 11, 418 | 0.45 | 0.93 |
|  |  | Sex* Day | 11, 418 | 1.04 | 0.41 |
|  |  | Genotype*Sex*Day | 11, 418 | 0.29 | 0.99 |
| **Eye Opening** | | | | | |
| 1st Day of occurrence | Y84X | Genotype | 1, 35.74 | 0.51 | 0.48 |
|  |  | Sex | 1, 33.82 | 0.04 | 0.84 |
|  |  | Genotype* Sex | 1, 34.62 | 1.59 | 0.22 |
|  | R1627X | Genotype | 1, 34.83 | 0.05 | 0.83 |
|  |  | Sex | 1, 33.39 | 0.04 | 0.85 |
|  |  | Genotype* Sex | 1, 34.52 | 0.02 | 0.90 |
| **Pinna Detachment** | | | | | |
| 1st Day of occurrence | Y84X | Genotype | 1, 33.97 | 0.01 | 0.95 |
|  |  | Sex | 1, 33.03 | 0.27 | 0.60 |
|  |  | Genotype* Sex | 1, 33.37 | 0.04 | 0.85 |
|  | R1627X | Genotype | 1, 32.80 | 1.28 | 0.27 |
|  |  | Sex | 1, 32.31 | 1.86 | 0.18 |
|  |  | Genotype* Sex^a^ | 1, 32.71 | 5.19 | **0.03** |

Linear-mixed effects model results for effects of genotype, sex, and day (including interactions) on early postnatal body weight (weight ~ genotype*sex*session + (1|litter/ID)), eye opening (eye day ~ genotype*sex + (1|litter)), and pinna detachment (pinna day ~ genotype*sex + (1|Litter)). Bold values indicate statistically significant effects (*p* ≤ 0.05). Post hoc comparisons were conducted following significant interactions using estimated marginal means.

^a^Post hoc comparisons were conducted to follow-up the significant genotype × sex interaction in *Scn2a^R1627X/+^* mice compared to wildtype for the day of occurrence for pinna detachment. We observed a significant difference between female wildtype and *Scn2a^R1627X/+^* mice (females, *t*(9.07) = -2.09, *p* = 0.05).

**Supplemental Table 2. Effects of the Y84X and R1627X variants on cliff avoidance**

| **Measure** | **Mouse Line** | **Effect** | **NumDF,DenDF** | **F-Value** | **P-Value** |
| --- | --- | --- | --- | --- | --- |
| Score | Y84X | Genotype | 1, 38.52 | 1.49 | 0.23 |
|  |  | Sex | 1, 35.09 | 1.34 | 0.26 |
|  |  | Day | 8, 311.25 | 69.45 | **<0.001** |
|  |  | Genotype* Sex | 1, 37.37 | 2.70 | 0.11 |
|  |  | Genotype* Day | 8, 311.25 | 0.85 | 0.56 |
|  |  | Sex* Day | 8, 311.25 | 0.47 | 0.88 |
|  |  | Genotype*Sex*Day | 8, 311.25 | 1.66 | 0.12 |
|  | R1627X | Genotype | 1, 310.13 | 0.23 | 0.63 |
|  |  | Sex | 1, 340.68 | 0.006 | 0.94 |
|  |  | Day | 8, 335.51 | 54.14 | **<0.001** |
|  |  | Genotype* Sex | 1, 320.57 | 0.0002 | 0.99 |
|  |  | Genotype* Day | 8, 335.51 | 0.54 | 0.83 |
|  |  | Sex* Day | 8, 335.51 | 1.70 | 0.10 |
|  |  | Genotype*Sex*Day | 8, 335.51 | 0.17 | 1 |
| Latency to Turn | Y84X | Genotype^a^ | 1, 37.24 | 6.84 | **0.01** |
|  |  | Sex | 1, 34.55 | 2.36 | 0.12 |
|  |  | Day | 8, 311.22 | 64.23 | **<0.001** |
|  |  | Genotype* Sex | 1, 35.94 | 0.88 | 0.35 |
|  |  | Genotype* Day^b^ | 8, 311.22 | 2.93 | **0.004** |
|  |  | Sex* Day | 8, 311.22 | 1 | 0.44 |
|  |  | Genotype*Sex*Day | 8, 311.22 | 0.92 | 0.50 |
|  | R1627X | Genotype | 1, 37.42 | 0.19 | 0.67 |
|  |  | Sex | 1, 34.38 | 0.35 | 0.56 |
|  |  | Day | 8, 304 | 48.18 | **<0.001** |
|  |  | Genotype* Sex | 1, 36.75 | 0.84 | 0.37 |
|  |  | Genotype* Day | 8, 304 | 0.23 | 0.99 |
|  |  | Sex* Day | 8, 304 | 1.40 | 0.200 |
|  |  | Genotype*Sex*Day | 8, 304 | 0.22 | 0.99 |
| 1st Day of occurrence | Y84X | Genotype | 1, 38.99 | 1.64 | 0.21 |
|  |  | Sex | 1, 36.64 | 0.96 | 0.33 |
|  |  | Genotype* Sex | 1, 38.81 | 1.97 | 0.17 |
|  | R1627X | Genotype | 1, 37.58 | 0.05 | 0.82 |
|  |  | Sex | 1, 34.74 | 0.28 | 0.60 |
|  |  | Genotype* Sex | 1, 37.24 | 0.07 | 0.80 |

Linear-mixed effects model results for effects of genotype, sex and their interactions for the performance (cliff avoidance measure ~ genotype*sex*session + (1|litter/ID)) and occurrence (cliff avoidance day ~ genotype*sex + (1|litter)) of cliff avoidance. Bold values indicate statistically significant effects (*p* ≤ 0.05). Post hoc comparisons were conducted following significant interactions using estimated marginal means.

^a^There was a significant main effect of genotype between wildtype and *Scn2a^Y84X/+^* mice for the latency to turn on the cliff avoidance assay (wildtype v. Y84X, F(D_1_,D_37.24_) = 6.84, *p* = 0.01). *Scn2a^Y84X/+^* mice were faster to turn than their wildtype littermates.

^b^Post hoc comparisons were conducted to follow-up the significant genotype × day interaction within the Y84X line for the latency to turn on the cliff avoidance assay. We observed a significant difference between wildtype and *Scn2a^Y84X/+^* mice on days 8 and 9 (PND 8, t(340) = 2.62, *p*= 0.01); (PND 9, t(340) = 4.62, *p* =<.0001).

**Supplemental Table 3. Effects of the Y84X and R1627X variants on the surface righting reflex**

| **Measure** | **Mouse Line** | **Effect** | **NumDF,DenDF** | **F-Value** | **P-Value** |
| --- | --- | --- | --- | --- | --- |
| Score | Y84X | Genotype | 1, 39 | 1.84 | 0.18 |
|  |  | Sex | 1, 39 | 1.72 | 0.20 |
|  |  | Day | 4, 156 | 20.05 | **<0.001** |
|  |  | Genotype* Sex^a^ | 1, 39 | 4.33 | **0.04** |
|  |  | Genotype* Day | 4, 156 | 1.04 | 0.39 |
|  |  | Sex* Day | 4, 156 | 0.44 | 0.78 |
|  |  | Genotype*Sex*Day | 4, 156 | 0.45 | 0.77 |
|  | R1627X | Genotype | 1, 35.05 | 0.12 | 0.73 |
|  |  | Sex | 1, 33.06 | 1.43 | 0.24 |
|  |  | Day | 5, 190 | 37.11 | **<0.001** |
|  |  | Genotype* Sex | 1, 34.61 | 1.01 | 0.32 |
|  |  | Genotype* Day | 5, 190 | 1.48 | 0.20 |
|  |  | Sex* Day | 5, 190 | 0.32 | 0.90 |
|  |  | Genotype*Sex*Day | 5, 190 | 1.67 | 0.14 |
| Latency to Right | Y84X | Genotype ^b^ | 1, 39 | 4.79 | **0.04** |
|  |  | Sex ^c^ | 1, 39 | 4.58 | **0.04** |
|  |  | Day | 4, 156 | 40.44 | **<0.001** |
|  |  | Genotype* Sex ^d^ | 1, 39 | 6.47 | **0.02** |
|  |  | Genotype* Day | 4, 156 | 1.55 | 0.19 |
|  |  | Sex* Day | 4, 156 | 0.53 | 0.71 |
|  |  | Genotype*Sex*Day | 4, 156 | 1.47 | 0.21 |
|  | R1627X | Genotype | 1, 227.32 | 0.28 | 0.60 |
|  |  | Sex | 1, 226.92 | 1 | 0.32 |
|  |  | Day | 5, 221.63 | 71.99 | **<0.001** |
|  |  | Genotype* Sex | 1, 227.82 | 0.11 | 0.75 |
|  |  | Genotype* Day | 5, 221.63 | 1.68 | 0.14 |
|  |  | Sex* Day | 5, 221.63 | 0.82 | 0.54 |
|  |  | Genotype*Sex*Day | 5, 221.63 | 0.62 | 0.68 |
| 1st Day of occurrence | Y84X | Genotype | 1, 38.98 | 1.02 | 0.32 |
|  |  | Sex | 1, 36.45 | 0.81 | 0.37 |
|  |  | Genotype* Sex | 1, 38.84 | 2.66 | 0.11 |
|  | R1627X | Genotype | 1, 37.83 | 1.22 | 0.28 |
|  |  | Sex | 1, 35.16 | 0.64 | 0.43 |
|  |  | Genotype* Sex | 1, 36.10 | 2.17 | 0.15 |

Linear-mixed effects model results for effects of genotype, sex and their interactions on performance (surface righting measure ~ genotype*sex*session + (1|litter/ID)) and occurrence (surface right day ~ genotype*sex + (1|litter)) of the surface righting reflex. Bold values indicate statistically significant effects (*p* ≤ 0.05). Post hoc comparisons were conducted following significant interactions using estimated marginal means.

^a^Post hoc comparisons were conducted to follow-up the significant genotype × sex interaction effect between wildtype and *Scn2a^Y84X/+^* mice for score on the surface righting reflex. We observed a significant difference between male wildtype and *Scn2a^Y84X/+^* mice (males, t(37.20) = -2.13, *p* = 0.04).

^b^There was a main effect of genotype between wildtype and *Scn2a^Y84X/+^* mice for the latency to right on the surface righting reflex (wildtype v. Y84X, F(D_1_,D_39_) = 4.79, *p* = 0.04). *Scn2a^Y84X/+^* mice were faster than their wildtype littermates.

^c^There was a significant main effect of sex between wildtype and *Scn2a^Y84X/+^* mice for the latency to right on the surface righting reflex (sex, F(D_1_,D_39_) = 4.58, *p* = 0.04). Male mice were faster than female mice.

^d^Post hoc comparisons were conducted to follow-up the significant genotype × sex interaction between *Scn2a^Y84X/+^* mice and wildtype littermates for the latency to right on the surface righting reflex. We observed a significant difference between male wildtype and *Scn2a^Y84X/+^* mice (males, t(37.20) = 2.93, *p* = 0.006).

**Supplemental Table 4. Effects of the Y84X and R1627X variants on negative geotaxis**

| **Measure** | **Mouse Line** | **Effect** | **NumDF,DenDF** | **F-Value** | **P-Value** |
| --- | --- | --- | --- | --- | --- |
| Score | Y84X | Genotype | 1, 36.19 | 0.44 | 0.51 |
|  |  | Sex | 1, 32.68 | 2.72 | 0.11 |
|  |  | Day | 6, 234 | 74.73 | **<0.001** |
|  |  | Genotype* Sex | 1, 34.34 | 0.45 | 0.59 |
|  |  | Genotype* Day | 6, 234 | 0.49 | 0.81 |
|  |  | Sex* Day | 6, 234 | 1.38 | 0.22 |
|  |  | Genotype*Sex*Day | 6, 234 | 0.96 | 0.45 |
|  | R1627X | Genotype | 1, 35.23 | 0.002 | 0.96 |
|  |  | Sex | 1, 33.42 | 1.712 | 0.20 |
|  |  | Day | 5, 190 | 55.33 | **<0.001** |
|  |  | Genotype* Sex | 1, 34.83 | 0.15 | 0.71 |
|  |  | Genotype* Day | 5, 190 | 1.33 | 0.25 |
|  |  | Sex* Day | 5, 190 | 1.18 | 0.32 |
|  |  | Genotype*Sex*Day | 5, 190 | 0.98 | 0.43 |
| Latency to Turn | Y84X | Genotype | 1, 36.13 | 1.11 | 0.30 |
|  |  | Sex | 1, 33.08 | 1.96 | 0.18 |
|  |  | Day | 9, 351 | 115.62 | **<0.001** |
|  |  | Genotype* Sex | 1, 34.48 | 1.44 | 0.24 |
|  |  | Genotype* Day | 9, 351 | 0.45 | 0.91 |
|  |  | Sex* Day | 9, 351 | 0.67 | 0.73 |
|  |  | Genotype*Sex*Day | 9, 351 | 1.25 | 0.26 |
|  | R1627X | Genotype | 1, 37.26 | 1 | 0.32 |
|  |  | Sex | 1, 33.42 | 2.62 | 0.12 |
|  |  | Day | 5, 190 | 8.36 | **<0.001** |
|  |  | Genotype* Sex | 1, 36.39 | 1.95 | 0.17 |
|  |  | Genotype* Day | 5, 190 | 0.47 | 0.80 |
|  |  | Sex* Day | 5, 190 | 1.08 | 0.37 |
|  |  | Genotype*Sex*Day | 5, 190 | 2.08 | 0.07 |
| 1st Day of occurrence | Y84X | Genotype | 1, 37.8 | 0.16 | 0.69 |
|  |  | Sex | 1, 33.61 | 0.25 | 0.62 |
|  |  | Genotype* Sex | 1, 36.07 | 0.14 | 0.71 |
|  | R1627X | Genotype | 1, 35.61 | 0.97 | 0.33 |
|  |  | Sex | 1, 34.20 | 1.28 | 0.27 |
|  |  | Genotype* Sex | 1, 34.63 | 0.89 | 0.35 |

Linear-mixed effects model results for effects of genotype, sex and their interactions for the performance (negative geotaxis measure ~ genotype*sex*session + (1|litter/ID)) and occurrence (negative geotaxis day ~ genotype*sex + (1|litter)) of negative geotaxis. Bold values indicate statistically significant effects (*p* ≤ 0.05).

**Supplemental Table 5. Effects of the Y84X and R1627X variants on the forelimb grasping reflex**

| **Measure** | **Mouse Line** | **Effect** | **NumDF,DenDF** | **F-Value** | ***P*-Value** |
| --- | --- | --- | --- | --- | --- |
| Score | Y84X | Genotype | 1, 34.86 | 0.69 | 0.41 |
|  |  | Sex | 1, 33.39 | 1.22 | 0.28 |
|  |  | Day | 7, 23 | 73.20 | **<0.001** |
|  |  | Genotype* Sex | 1, 33.96 | 0.09 | 0.77 |
|  |  | Genotype* Day | 7, 234 | 1.52 | 0.17 |
|  |  | Sex* Day | 7, 234 | 1.44 | 0.20 |
|  |  | Genotype*Sex*Day | 7, 234 | 1.06 | 0.39 |
|  | R1627X | Genotype | 1, 33.01 | 0.003 | 0.96 |
|  |  | Sex | 1, 32.48 | 0.02 | 0.89 |
|  |  | Day | 4, 151 | 88.53 | **<0.001** |
|  |  | Genotype* Sex | 1, 33.13 | 3.08 | 0.09 |
|  |  | Genotype* Day | 4, 151 | 1.36 | 0.25 |
|  |  | Sex* Day | 4, 151 | 0.88 | 0.48 |
|  |  | Genotype*Sex*Day | 4, 151 | 2.10 | 0.08 |
| 1st Day of occurrence | Y84X | Genotype | 1, 38.98 | 0.58 | 0.45 |
|  |  | Sex | 1, 36.45 | 0.57 | 0.46 |
|  |  | Genotype* Sex | 1, 38.84 | 2.51 | 0.12 |
|  | R162  7X | Genotype | 1, 33.51 | 0.02 | 0.88 |
|  |  | Sex | 1, 32.60 | 0.001 | 0.98 |
|  |  | Genotype* Sex | 1, 33.33 | 2.32 | 0.14 |

Linear-mixed effects model results for effects of genotype, sex and their interactions on performance and occurrence of the forepaw grasping reflex (grasp ~ genotype*sex*session + (1|litter/ID)). Bold values indicate statistically significant effects (*p* ≤ 0.05).

**Supplemental Table 6. Effects of the Y84X and R1627X variants on ultrasonic vocalizations (USV)**

| **Measure** | **Mouse Line** | **Effect** | **NumDF,DenDF** | **F-Value** | ***P*-Value** |
| --- | --- | --- | --- | --- | --- |
| Count | Y84X | Genotype | 1, 40.32 | 0.02 | 0.89 |
|  |  | Sex | 1, 38.75 | 2.59 | 0.12 |
|  |  | Genotype* Sex | 1, 38.49 | 1.02 | 0.32 |
|  | R1627X | Genotype | 1, 33.30 | 0.02 | 0.89 |
|  |  | Sex | 1, 35.03 | 0.21 | 0.65 |
|  |  | Genotype* Sex | 1, 34.46 | 1.24 | 0.27 |
| Average Duration | Y84X | Genotype | 1, 40.99 | 0.79 | 0.38 |
|  |  | Sex | 1, 39.23 | 0.62 | 0.44 |
|  |  | Genotype* Sex | 1, 38.88 | 0.03 | 0.86 |
|  | R1627X | Genotype | 1, 36 | 0.27 | 0.61 |
|  |  | Sex | 1, 36 | 0.71 | 0.40 |
|  |  | Genotype* Sex | 1, 36 | 0.23 | 0.64 |
| Mean Frequency | Y84X | Genotype | 1, 42.44 | 1.24 | 0.27 |
|  |  | Sex | 1, 42.88 | 1.07 | 0.31 |
|  |  | Genotype* Sex | 1, 42.24 | 0.33 | 0.57 |
|  | R1627X | Genotype | 1, 36 | 0.49 | 0.49 |
|  |  | Sex | 1, 36 | 0.79 | 0.38 |
|  |  | Genotype* Sex | 1, 36 | 1 | 0.32 |
| Average Number of Modulations | Y84X | Genotype | 1, 42.94 | 0.77 | 0.39 |
|  |  | Sex | 1, 42.38 | 0.44 | 0.51 |
|  |  | Genotype* Sex | 1, 41.60 | 0.88 | 0.36 |
|  | R1627X | Genotype | 1, 34.31 | 0.95 | 0.34 |
|  |  | Sex | 1, 35.63 | 1.35 | 0.25 |
|  |  | Genotype* Sex | 1, 35.20 | 2.85 | 0.10 |

Linear-mixed effects model results for effects of genotype, sex and their interactions on ultrasonic vocalizations (USV metric ~ genotype*sex + (1|litter)). Bold values indicate statistically significant effects (*p* ≤ 0.05).

**Supplemental Table 7. Y84X and R1627X variant effects on the prevalence of Vglut1 cerebellar puncta in mice**

| **Age** | **Mouse Line** | **Effect** | **NumDF,DenDF** | **F-Value** | **P-Value** |
| --- | --- | --- | --- | --- | --- |
| PND6 | Y84X | Genotype | 1, 10.78 | 2.54 | 0.14 |
|  |  | Lobule | 7, 68.40 | 1.50 | 0.18 |
|  |  | Sex | 1, 10.78 | 3.47 | 0.09 |
|  |  | Genotype* Lobule | 7, 68.40 | 0.19 | 0.99 |
|  |  | Genotype* Sex | 1, 10.78 | 0.48 | 0.51 |
|  |  | Sex* Lobule | 7, 68.40 | 0.89 | 0.52 |
|  |  | Genotype*Sex*Lobule | 7, 68.40 | 1.32 | 0.26 |
|  | R1627X | Genotype | 1, 10.07 | 1.69 | 0.22 |
|  |  | Lobule | 7, 66.57 | 6.32 | **<0.001** |
|  |  | Sex^a^ | 1, 10.07 | 5.34 | **0.04** |
|  |  | Genotype* Lobule | 7, 66.57 | 0.65 | 0.71 |
|  |  | Genotype* Sex | 1, 10.07 | 1.83 | 0.21 |
|  |  | Sex* Lobule | 7, 66.57 | 1.42 | 0.21 |
|  |  | Genotype*Sex*Lobule | 7, 66.57 | 0.69 | 0.68 |
| PND12 | Y84X | Genotype^b^ | 1, 12.46 | 6.89 | **0.02** |
|  |  | Lobule | 7, 82.44 | 5.53 | **<0.001** |
|  |  | Sex | 1, 12.46 | 0.34 | 0.57 |
|  |  | Genotype* Lobule^c^ | 7, 82.44 | 2.21 | **0.04** |
|  |  | Genotype* Sex | 1, 12.46 | 1.23 | 0.29 |
|  |  | Sex* Lobule | 7, 82.44 | 1 | 0.44 |
|  |  | Genotype*Sex*Lobule | 7, 82.44 | 0.84 | 0.56 |
|  | R1627X | Genotype | 1, 8.69 | 3.28 | 0.11 |
|  |  | Lobule | 7, 61.21 | 2.72 | **<0.05** |
|  |  | Sex | 1, 8.14 | 0.16 | 0.70 |
|  |  | Genotype* Lobule | 7, 61.21 | 1.09 | 0.38 |
|  |  | Genotype* Sex | 1, 8.69 | 0.02 | 0.90 |
|  |  | Sex* Lobule | 7, 61.21 | 1.47 | 0.19 |
|  |  | Genotype*Sex*Lobule^d^ | 7, 61.21 | 2.53 | **0.02** |
| PND21 | Y84X | Genotype | 1, 10.08 | 2.70 | 0.13 |
|  |  | Lobule | 7, 67.30 | 3.76 | **<0.001** |
|  |  | Sex | 1, 10.17 | 2.47 | 0.15 |
|  |  | Genotype* Lobule | 7, 67.30 | 0.27 | 0.96 |
|  |  | Genotype* Sex | 1, 6.65 | 0.12 | 0.74 |
|  |  | Sex* Lobule | 7, 67.30 | 0.39 | 0.90 |
|  |  | Genotype*Sex*Lobule | 7, 67.30 | 1.41 | 0.22 |
|  | R1627X | Genotype | 1, 5.57 | 0.003 | 0.96 |
|  |  | Lobule | 7, 63 | 10.52 | **<0.001** |
|  |  | Sex | 1, 7.67 | 0.33 | 0.58 |
|  |  | Genotype* Lobule | 7, 63 | 0.41 | 0.89 |
|  |  | Genotype* Sex | 1, 1.15 | 0.02 | 0.92 |
|  |  | Sex* Lobule | 7, 63 | 0.57 | 0.78 |
|  |  | Genotype*Sex*Lobule | 7, 63 | 0.66 | 0.71 |

Linear-mixed effects model results for effects of genotype, sex and their interactions on the prevalence of Vglut1 cerebellar puncta (Vglut1 coverage ~ genotype*sex*lobule + (1|litter/ID)). Bold values indicate statistically significant effects (*p* ≤ 0.05). Post hoc comparisons were conducted following significant interactions using estimated marginal means.

^a^There was a significant main effect of sex between wildtype and *Scn2a^R1627X/+^* mice at PND6 for Vglut1 cerebellar puncta expression (wildtype v. R1627X, sex, F(D_1_,D_10.07_) = 5.34, *p* = 0.04). Males had increased expression compared to females.

^b^There was a significant main effect of genotype between wildtype and *Scn2a^Y84X/+^* mice at PND12 for Vglut1 cerebellar puncta expression (wildtype v. Y84X, F(D_1_,D_12.46_) = 6.89, *p* = 0.02). *Scn2a^Y84X/+^* mice had decreased Vglut1 expression compared to wildtype littermates.

^c^Post hoc comparisons were conducted to follow-up the significant genotype × lobule interaction effect between *Scn2a^Y84X/+^* mice and wildtype littermates at PND12 for Vglut1 cerebellar puncta expression. We observed a significant difference between wildtype and *Scn2a^Y84X/+^* mice in cerebellar lobules IV/V and VII (lobule IV/V, t(72.40) = 3.33, *p* = 0.001); (lobule VII, t(73.70) = 3.38, *p* = 0.001).

^d^Post hoc comparisons were conducted to follow-up the significant interaction effect genotype × sex × lobule between *Scn2a^R1627X/+^* mice and wildtype littermates at PND12 for Vglut 1 cerebellar puncta expression. We observed a significant difference between male wildtype and *Scn2a^R1627X/+^* mice in cerebellar lobules VIII and IX (lobule VIII males, ΔM = -2.87, t(53.50) = -2.52, *p* = 0.02) ;(Lobule IX males, t(53.50) = -2.86, *p* = 0.006).

**Supplemental Table 8. Y84X and R1627X variant effects on the prevalence of Vglut2 cerebellar puncta in mice**

| **Age** | **Mouse Line** | **Effect** | **NumDF,DenDF** | **F-Value** | **P-Value** |
| --- | --- | --- | --- | --- | --- |
| PND21 | Y84X | Genotype^a^ | 1, 11.04 | 19.24 | **0.001** |
|  |  | Lobule | 7, 72.23 | 1.49 | 0.18 |
|  |  | Sex | 1, 11.04 | 0.44 | 0.52 |
|  |  | Genotype* Lobule | 7, 72.23 | 1.01 | 0.43 |
|  |  | Genotype* Sex | 1, 11.04 | 1.80 | 0.21 |
|  |  | Sex* Lobule | 7, 72.23 | 1.83 | 0.10 |
|  |  | Genotype*Sex*Lobule | 7, 72.23 | 1.06 | 0.40 |
|  | R1627X | Genotype | 1, 8.77 | 0.20 | 0.67 |
|  |  | Lobule | 7, 58.99 | 8.12 | **<0.001** |
|  |  | Sex | 1, 8.77 | 0.06 | 0.82 |
|  |  | Genotype* Lobule | 7, 58.99 | 1.61 | 0.15 |
|  |  | Genotype* Sex | 1, 8.77 | 0.32 | 0.59 |
|  |  | Sex* Lobule | 7, 58.99 | 0.97 | 0.46 |
|  |  | Genotype*Sex*Lobule | 7, 58.99 | 0.89 | 0.52 |

Linear-mixed effects model results for effects of genotype, sex and their interactions on the prevalence of Vglut2 cerebellar puncta (Vglut2 coverage ~ genotype*sex*lobule + (1|litter/ID)). Bold values indicate statistically significant effects (*p* ≤ 0.05). Post hoc comparisons were conducted following significant interactions using estimated marginal means.

^a^There was a significant main effect of genotype between *Scn2a^Y84X/+^* mice and wildtype littermates at PND21 for the prevalence of Vglut2 positive cerebellar puncta (wildtype v. Y84X, F(D_1_,D_11.04_) = 19.24, *p* = 0.001). *Scn2a^Y84X/+^* mice had increased Vglut2 puncta compared to wildtype littermates.

**Supplemental Table 9. Y84X and R1627X variant effects on the cerebellar protein expression of Vglut1 and Vglut2**

| **Measure** | **Mouse Line** | **Effect** | **NumDF,DenDF** | **F-Value** | **P-Value** |
| --- | --- | --- | --- | --- | --- |
| Vglut1 | Y84X | Genotype^a^ | 1,7.58 | 5.71 | **0.05** |
|  |  | Sex | 1, 6.51 | 3.76 | 0.10 |
|  |  | Genotype* Sex | 1, 7.86 | 0.25 | 0.63 |
|  | R1627X | Genotype | 1, 7.98 | 1.66 | 0.23 |
|  |  | Sex | 1, 6.76 | 4.12 | 0.08 |
|  |  | Genotype* Sex | 1, 6.76 | 2.95 | 0.13 |
| Vglut2 | Y84X | Genotype^b^ | 1, 4.77 | 14.05 | **0.02** |
|  |  | Sex^c^ | 1, 4.61 | 25.21 | **0.01** |
|  |  | Genotype* Sex^d^ | 1, 5.04 | 16.92 | **0.01** |
|  | R1627X | Genotype^e^ | 1, 7.91 | 7.20 | **0.03** |
|  |  | Sex | 1, 6.86 | 3.21 | 0.12 |
|  |  | Genotype* Sex | 1, 6.86 | 0.29 | 0.61 |

Linear-mixed effects model results for effects of genotype, sex and their interactions on the prevalence of Vglut1 and Vglut2 cerebellar protein expression (Vglut1 or Vglut2 ~ genotype*sex + (1|litter)). Bold values indicate statistically significant effects (*p* ≤ 0.05). Post hoc comparisons were conducted following significant interactions using estimated marginal means.

^a^There was a significant main effect of genotype effect between wildtype and *Scn2a^Y84X/+^* mice for Vglut1 cerebellar protein expression. We observed decreased Vglut1 cerebellar protein expression between *Scn2a^Y84X/+^* mice and wildtype littermates at PND21 (wildtype v. Y84X, F(D_1_,D_7.58_) = 5.71, *p* = 0.05).

^b^We observed a significant main effect of genotype between *Scn2a^Y84X/+^* mice and wildtype littermates at PND21 for Vglut2 positive puncta (wildtype v. Y84X, F(D_1_,D_4.77_) = 14.05, *p* = 0.02). *Scn2a^Y84X/+^* mice had increased Vglut2 protein expression compared to wildtype littermates.

^c^There was a significant main effect of sex between wildtype and *Scn2a^Y84X/+^* mice at PND21 for cerebellar Vglut2 protein expression (sex, F(D_1_,D_4.77_) = 14.05, *p* = 0.02). Females had greater Vglut2 protein expression than males.

^d^Post hoc comparisons were conducted to follow-up the significant genotype x sex interaction effect between *Scn2a^Y84X/+^* mice and wildtype littermates for Vglut2 cerebellar protein expression. We found a significant difference between male wildtype and *Scn2a^Y84X/+^* mice (males, t(5.81) = -5.21, *p* = 0.002).

**^e^**There was a significant main effect of genotype between wildtype and *Scn2a^R1627X/+^* mice for Vglut2 cerebellar protein expression (wildtype v. R1627X, F(D_1_,D_7.91_) = 7.20, *p* = 0.03). *Scn2a^R1627X/+^* mice had decreased Vglut2 protein expression compared to wildtype littermates.

**Supplemental Table 10. Effects of the Y84X and R1627X variants on Purkinje Cell Density**

| **Age** | **Mouse Line** | **Effect** | **NumDF,DenDF** | **F-Value** | **P-Value** |
| --- | --- | --- | --- | --- | --- |
| PND12 | Y84X | Genotype^a^ | 1, 11.16 | 5.75 | **0.04** |
|  |  | Lobule | 7, 75.40 | 5.95 | **<0.001** |
|  |  | Sex | 1, 11.16 | 0.06 | 0.81 |
|  |  | Genotype* Lobule | 7, 75.40 | 0.95 | 0.47 |
|  |  | Genotype* Sex | 1, 11.16 | 0.29 | 0.60 |
|  |  | Sex* Lobule | 7, 75.40 | 0.89 | 0.52 |
|  |  | Genotype*Sex*Lobule | 7, 75.40 | 1.73 | 0.12 |
|  | R1627X | Genotype | 1, 8.06 | 0.06 | 0.82 |
|  |  | Lobule | 7, 55.12 | 7.72 | **<0.001** |
|  |  | Sex | 1, 8.06 | 0.07 | 0.81 |
|  |  | Genotype* Lobule | 7, 55.12 | 1.54 | 0.17 |
|  |  | Genotype* Sex | 1, 8.06 | 1.46 | 0.26 |
|  |  | Sex* Lobule | 7, 55.12 | 0.24 | 0.97 |
|  |  | Genotype*Sex*Lobule^b^ | 7, 55.12 | 2.16 | **0.05** |
| PND21 | Y84X | Genotype^c^ | 1, 72 | 21.27 | **<0.001** |
|  |  | Lobule | 7, 72 | 15.85 | **<0.001** |
|  |  | Sex | 1, 72 | 0.41 | 0.53 |
|  |  | Genotype* Lobule | 7, 72 | 0.86 | 0.54 |
|  |  | Genotype* Sex | 1, 72 | 1.05 | 0.31 |
|  |  | Sex* Lobule | 7, 72 | 1.16 | 0.34 |
|  |  | Genotype*Sex*Lobule | 7, 72 | 1.16 | 0.34 |
|  | R1627X | Genotype^d^ | 1, 6.69 | 10.53 | **0.02** |
|  |  | Lobule | 7, 56 | 4.68 | **<0.001** |
|  |  | Sex | 1, 7.06 | 0.03 | 0.86 |
|  |  | Genotype* Lobule | 7, 56 | 0.75 | 0.63 |
|  |  | Genotype* Sex | 1, 4.99 | 0.97 | 0.37 |
|  |  | Sex* Lobule | 7, 56 | 1.37 | 0.24 |
|  |  | Genotype*Sex*Lobule | 7, 56 | 1.23 | 0.30 |

Linear-mixed effects model results for effects of genotype, sex and their interactions on Purkinje cell density (Purkinje density ~ genotype*sex*lobule + (1|litter/ID)). Bold values indicate statistically significant effects (*p* ≤ 0.05). Post hoc comparisons were conducted following significant interactions using estimated marginal means.

^a^There was a significant main effect of genotype between wildtype and *Scn2a^Y84X/+^* mice at PND12 for Purkinje cell density (wildtype v. Y84X, F(D_1_,D_11.16_) = 5.75, *p* = 0.04). *Scn2a^Y84X/+^* mice at PND12 had decreased Purkinje cell density compared to wildtype littermates.

^b^Post hoc comparisons were conducted to follow-up the significant genotype × sex × lobule interaction between wildtype and *Scn2a^R1627X/+^* mice at PND12 for Purkinje cell density. We observed a significant difference between female wildtype and *Scn2a^R1627X/+^* mice in cerebellar lobule III (lobule III, females, t(42.50) = -2.61, *p* = 0.01) and a strong trend in cerebellar lobule VIII (lobule VIII females, t(42.5) = -1.93, *p* = 0.06).

^c^We observed a significant main effect of genotype between wildtype and *Scn2a^Y84X/+^* mice at PND21 for Purkinje cell density (wildtype v. Y84X, F(D_1_,D_72_) = 21.27, *p* = <0.001). *Scn2a^Y84X/+^* mice at PND21 had decreased Purkinje cell density compared to wildtype littermates.

^d^There was a significant main effect of genotype between wildtype and *Scn2a^R1627X/+^* mice at PND21 for Purkinje cell density (wildtype v. R1627X, F(D_1_,D_6.69_) = 10.53, *p* = 0.02). *Scn2a^R1627X/+^* mice at PND21 had decreased Purkinje cell density compared to wildtype littermates.

**Supplemental Table 11. Effects of the Y84X and R1627X variants on Purkinje Cell soma area**

| **Age** | **Mouse Line** | **Effect** | **NumDF,DenDF** | **F-Value** | **P-Value** |
| --- | --- | --- | --- | --- | --- |
| PND12 | Y84X | Genotype^a^ | 1, 7.25 | 10.72 | **0.01** |
|  |  | Lobule | 7, 77 | 6.32 | **<0.001** |
|  |  | Sex^b^ | 1, 9.91 | 5.21 | **0.05** |
|  |  | Genotype* Lobule | 7, 77 | 1.41 | 0.22 |
|  |  | Genotype* Sex | 1, 9.90 | 0.72 | 0.42 |
|  |  | Sex* Lobule | 7, 77 | 1.60 | 0.15 |
|  |  | Genotype*Sex*Lobule | 7, 77 | 0.68 | 0.69 |
|  | R1627X | Genotype | 1, 7.20 | 0.22 | 0.65 |
|  |  | Lobule | 7, 56 | 10.34 | **<0.001** |
|  |  | Sex | 1, 7 | 2.81 | 0.14 |
|  |  | Genotype* Lobule | 7, 56 | 0.76 | 0.63 |
|  |  | Genotype* Sex | 1, 7.20 | 0.87 | 0.38 |
|  |  | Sex* Lobule | 7, 56 | 1.16 | 0.34 |
|  |  | Genotype*Sex*Lobule | 7, 56 | 1.44 | 0.21 |
| PND21 | Y84X | Genotype^c^ | 1, 8.30 | 6.96 | **0.03** |
|  |  | Lobule | 7, 63 | 16.96 | **<0.001** |
|  |  | Sex | 1, 4.69 | 0.10 | 0.77 |
|  |  | Genotype* Lobule | 7, 63 | 1.13 | 0.36 |
|  |  | Genotype* Sex | 1, 4.82 | 0.23 | 0.65 |
|  |  | Sex* Lobule | 7, 63 | 1 | 0.44 |
|  |  | Genotype*Sex*Lobule^d^ | 7, 63 | 2.82 | **0.01** |
|  | R1627X | Genotype^e^ | 1, 6.34 | 6.59 | **0.04** |
|  |  | Lobule | 7, 56 | 9.88 | **<0.001** |
|  |  | Sex^f^ | 1, 6.62 | 5.94 | **0.05** |
|  |  | Genotype* Lobule | 7, 56 | 1.75 | 0.12 |
|  |  | Genotype* Sex | 1, 6.99 | 0.88 | 0.38 |
|  |  | Sex* Lobule | 7, 56 | 1.43 | 0.21 |
|  |  | Genotype*Sex*Lobule | 7, 56 | 1.10 | 0.38 |

Linear-mixed effects model results for effects of genotype, sex and their interactions on Purkinje cell soma size (Purkinje soma ~ genotype*sex*lobule + (1|litter/ID)). Bold values indicate statistically significant effects (*p* ≤ 0.05). Post hoc comparisons were conducted following significant interactions using estimated marginal means.

^a^We observed a significant main effect of genotype between wildtype and *Scn2a^Y84X/+^* mice at PND12 for Purkinje cell soma size (wildtype v Y84X, F(D_1_,D_7.25_) = 10.72, *p* = 0.01). *Scn2a^Y84X/+^* mice at PND12 had decreased Purkinje cell soma area compared to wildtype littermates.

^b^There was a significant main effect of sex between wildtype and *Scn2a^Y84X/+^* mice at PND12 for Purkinje cell soma size (sex, F(D_1_,D_9.91_) = 5.21, *p* = 0.05). Males had smaller Purkinje cell somas than females.

^c^There was a significant main effect of genotype between wildtype and *Scn2a^Y84X/+^* mice at PND21 for Purkinje cell soma size (wildtype v Y84X, F(D_1_,D_8.30_) = 6.96, *p* = 0.03). *Scn2a^Y84X/+^* mice at PND21 had decreased Purkinje cell soma areas compared to wildtype littermates.

^d^Post hoc comparisons were conducted to follow-up the significant interaction effect between genotype × sex × lobule between wildtype and *Scn2a^Y84X/+^* mice at PND21 for Purkinje cell soma size. We observed a significant difference between female wildtype and *Scn2a^Y84X/+^* mice in cerebellar lobule II (lobule II, females, t(42.20) = 3.93, *p* = 0.0003) and male wildtype and *Scn2a^Y84X/+^* mice in lobule VI (lobule VI, males, t(51.70) = 3.51, *p* = 0.0009) as well as a trend for males in lobule VII (lobule VII, males, t(51.70) = 1.85, *p* = 0.07).

^e^There was a significant main effect of genotype between wildtype and *Scn2a^R1627X/+^* mice at PND21 for Purkinje cell soma size (wildtype v. R1627X, F(D_1_,D_6.34_) = 6.59, *p* = 0.04). *Scn2a^R1627X/+^* mice at PND21 had decreased Purkinje cell soma area compared to wildtype littermates.

^f^There was a significant main effect of sex between wildtype and *Scn2a^R1627X/+^* mice at PND21 for Purkinje cell soma size (sex, F(D_1_,D_6.62_) = 5.94, *p* = 0.05). Males had smaller Purkinje cell somas than females.

**Supplemental Table 12. Y84X and R1627X variant effects on external cerebellar granule layer thickness**

| **Age** | **Mouse Line** | **Effect** | **NumDF,DenDF** | **F-Value** | **P-Value** |
| --- | --- | --- | --- | --- | --- |
| PND6 | Y84X | Genotype | 1, 79.28 | 0.0001 | 0.99 |
|  |  | Lobule | 7, 78.39 | 14.64 | **<0.001** |
|  |  | Sex | 1, 83.79 | 0.66 | 0.44 |
|  |  | Genotype* Lobule | 7, 78.39 | 0.55 | 0.80 |
|  |  | Genotype* Sex^a^ | 1, 56.41 | 4.79 | **0.03** |
|  |  | Sex* Lobule | 7, 78.39 | 0.57 | 0.78 |
|  |  | Genotype*Sex*Lobule | 7, 78.39 | 1.25 | 0.29 |
|  | R1627X | Genotype | 1, 10.61 | 3.15 | 0.11 |
|  |  | Lobule | 7, 73.33 | 18.02 | **<0.001** |
|  |  | Sex | 1, 10 | 0.03 | 0.87 |
|  |  | Genotype* Lobule | 7, 73.33 | 1.47 | 0.19 |
|  |  | Genotype* Sex | 1, 11.20 | 0.004 | 0.95 |
|  |  | Sex* Lobule | 7, 73.33 | 1.97 | 0.08 |
|  |  | Genotype*Sex*Lobule^b^ | 7, 73.33 | 2.92 | **0.01** |
| PND12 | Y84X | Genotype | 1, 7.58 | 2.92 | 0.13 |
|  |  | Lobule | 7, 77 | 23.00 | **<0.001** |
|  |  | Sex | 1, 8.71 | 1.36 | 0.28 |
|  |  | Genotype* Lobule | 7, 77 | 1.04 | 0.41 |
|  |  | Genotype* Sex | 1, 8.66 | 3.62 | 0.09 |
|  |  | Sex* Lobule | 7, 77 | 0.80 | 0.59 |
|  |  | Genotype*Sex*Lobule^c^ | 7, 77 | 2.14 | **0.05** |
|  | R1627X | Genotype^d^ | 1, 7.03 | 9.07 | **0.02** |
|  |  | Lobule | 7, 56 | 19.35 | **<0.001** |
|  |  | Sex | 1, 7 | 0.05 | 0.83 |
|  |  | Genotype* Lobule | 7, 56 | 0.56 | 0.79 |
|  |  | Genotype* Sex^e^ | 1, 7.03 | 7.23 | **0.03** |
|  |  | Sex* Lobule^f^ | 7, 56 | 2.21 | **0.05** |
|  |  | Genotype*Sex*Lobule | 7, 56 | 1 | 0.44 |

Linear-mixed effects model results for effects of genotype, sex and their interactions on the 2- dimensional thickness of the external cerebellar granule layer (EGL) in mice (EGL ~ genotype*sex*lobule + (1|litter/ID)). Bold values indicate statistically significant effects (*p* ≤ 0.05). Post hoc comparisons were conducted following significant interactions using estimated marginal means.

^a^Post hoc comparisons were conducted to follow-up the significant interaction effect between genotype × sex between wildtype and *Scn2a^Y84X/+^* mice at PND6 in the external granule layer. We observed no difference between male and female mice when considering genotype (female, t(73.30) = -1.62, *p* = 0.11); (male, t(65.04) = 1.50, *p* =0.14).

^b^Post hoc comparisons were conducted to follow-up the significant interaction effect between genotype × sex × lobule between wildtype and *Scn2a^R1627X/+^* mice at PND6 in the external granule layer. We observed a strong trend between male wildtype and *Scn2a^R1627X/+^* mice in lobule II (lobule II, males, t(52.60) = 1.93, *p* = 0.06) in addition to significant difference in lobule III (lobule III, males, t(52.60) = 3.05, *p* = 0.004) and a significant difference between female wildtype and *Scn2a^R1627X/+^* mice in lobule VIII (lobule VIII, females, t(57.80) = 2.86, *p* = 0.01).

^c^Post hoc comparisons were conducted to follow-up the significant interaction effect between genotype × sex × lobule between wildtype and *Scn2a^Y84X/+^* mice at PND12 in the external granule layer. We observed a significant difference between female wildtype and *Scn2a^Y84X/+^* mice in cerebellar lobules II, III and IV/V (lobule II, females, t(29.70) = 2.99, *p* = 0.01); (lobule III, females, t(29.70) = 2.98, *p* = 0.01); (lobule IV/V, Female female, t(29.70) = 2.31, *p* = 0.03).

^d^There was a main effect of genotype between wildtype and *Scn2a^R1627X/+^* mice at PND12 in the external granule layer (wildtype v. R1627X, F(D_1_,D_7.03_) = 9.07, *p* = 0.02). *Scn2a^R1627X/+^* mice at PND12 had a thinner external granule layer compared to wildtype littermates.

**^e^**Post hoc comparisons were conducted to follow-up the significant interaction effect between genotype × sex between wildtype and *Scn2a^R1627X/+^* mice at PND12 in the external granule layer. We observed a significant difference between female wildtype and *Scn2a^R1627X/+^* mice (female, t(7.05) = 3.75, *p* = 0.01).

**^f^**Post hoc comparisons were conducted to follow-up the significant interaction effect between sex × lobule at PND12 between wildtype and *Scn2a^R1627X/+^* mice in the external granule layer. We observed a significant difference between male and female mice in cerebellar lobules VI (lobule IV, sex, t(61.60) = -2.52, *p* = 0.01).

**Supplemental Table 13. Y84X and R1627X variant effects on cerebellar molecular layer thickness**

| **Age** | **Mouse Line** | **Effect** | **NumDF,DenDF** | **F-Value** | **P-Value** |
| --- | --- | --- | --- | --- | --- |
| PND12 | Y84X | Genotype | 1, 8.20 | 0.60 | 0.46 |
|  |  | Lobule | 7, 77 | 10.53 | **<0.001** |
|  |  | Sex | 1, 9.77 | 0.87 | 0.37 |
|  |  | Genotype* Lobule | 7, 77 | 0.81 | 0.58 |
|  |  | Genotype* Sex | 1, 9.73 | 1.32 | 0.28 |
|  |  | Sex* Lobule | 7, 77 | 0.91 | 0.51 |
|  |  | Genotype*Sex*Lobule | 7, 77 | 1.64 | 0.14 |
|  | R1627X | Genotype | 1, 7.11 | 0.03 | 0.87 |
|  |  | Lobule | 7, 56 | 19.66 | **<0.001** |
|  |  | Sex | 1, 7 | 1.46 | 0.27 |
|  |  | Genotype* Lobule | 7, 56 | 0.83 | 0.57 |
|  |  | Genotype* Sex | 1, 7.11 | 1.55 | 0.25 |
|  |  | Sex* Lobule | 7, 56 | 1.42 | 0.22 |
|  |  | Genotype*Sex*Lobule | 7, 56 | 0.40 | 0.90 |
| PND21 | Y84X | Genotype | 1, 9 | 0.39 | 0.55 |
|  |  | Lobule | 7, 63 | 12.57 | **<0.001** |
|  |  | Sex | 1, 9 | 0.39 | 0.55 |
|  |  | Genotype* Lobule | 7, 63 | 0.28 | 0.96 |
|  |  | Genotype* Sex | 1, 9 | 2.53 | 0.15 |
|  |  | Sex* Lobule | 7, 63 | 1.52 | 0.18 |
|  |  | Genotype*Sex*Lobule | 7, 3 | 0.39 | 0.91 |
|  | R1627X | Genotype | 1, 23.83 | 0.02 | 0.90 |
|  |  | Lobule | 7, 57.15 | 13.48 | **<0.001** |
|  |  | Sex | 1, 20.90 | 1.27 | 0.27 |
|  |  | Genotype* Lobule | 7, 57.15 | 0.58 | 0.77 |
|  |  | Genotype* Sex | 1, 5 | 0.56 | 0.49 |
|  |  | Sex* Lobule | 7, 57.15 | 0.41 | 0.89 |
|  |  | Genotype*Sex*Lobule | 7, 57.15 | 0.30 | 0.95 |

Linear-mixed effects model results for effects of genotype, sex and their interactions on the 2- dimensional thickness of the cerebellar molecular layer (ML) in mice (ML ~ genotype*sex*lobule + (1|litter/ID)). Bold values indicate statistically significant effects (*p* ≤ 0.05).
